# Esm-1 mediates transcriptional polarization associated with diabetic kidney disease

**DOI:** 10.1101/2023.03.01.530562

**Authors:** Alexandre Gaudet, Xiaoyi Zheng, Neeraja Kambham, Vivek Bhalla

## Abstract

**Background:** Esm-1, endothelial cell-specific molecule-1, is a susceptibility gene for diabetic kidney disease (DKD) and is a cytokine- and glucose-regulated, secreted proteoglycan, that is notably expressed in kidney and attenuates inflammation and albuminuria. *Esm1* has restricted expression at the vascular tip during development but little is known about its expression pattern in mature tissues, and its precise effects in diabetes.

**Methods:** We utilized publicly available single-cell RNA sequencing data to explore the characteristics of *Esm1* expression in 27,786 renal endothelial cells obtained from four adult human and three mouse databases. We validated our findings using bulk transcriptome data from an additional 20 healthy subjects and 41 patients with DKD and using RNAscope. Using correlation matrices, we relate Esm1 expression to the glomerular transcriptome and evaluated these matrices with systemic over-expression of Esm-1.

**Results:** In both mice and humans, *Esm1* is expressed in a subset of all renal endothelial cell types and represents a minority of glomerular endothelial cells. In patients, *Esm1*(+) cells exhibit a highly conserved enrichment for blood vessel development genes. With diabetes, these cells are fewer in number and profoundly shift expression to reflect chemotaxis pathways. Analysis of these gene sets highlight candidate genes such as *Igfbp5* for cross talk between cell types. We also find that diabetes induces correlations in the expression of large clusters of genes, within cell type-enriched transcripts. *Esm1* significantly correlates with a majority genes within these clusters, delineating a glomerular transcriptional polarization reflected by the magnitude of *Esm1* deficiency. In diabetic mice, these gene clusters link *Esm1* expression to albuminuria, and over-expression of Esm-1 reverses the expression pattern in many of these genes.

**Conclusions:** A comprehensive analysis of single cell and bulk transcriptomes demonstrates that diabetes correlates with lower *Esm1* expression and with changes in the functional characterization of *Esm1*(+) cells. *Esm1* is both a marker for glomerular transcriptional polarization, and a mediator that re-orients the transcriptional program in DKD.

## Introduction

Diabetic kidney disease (DKD) is a common complication of diabetes mellitus, with a prevalence of ∼15% and is associated with poor long-term outcomes^1–3^. Chronic inflammation is a major contributor to kidney disease in diabetes^4–12^. Macrophage infiltration in the kidney correlates with the development of DKD^13–15^, while constitutive deletion of lymphocytes, cell adhesion and chemoattractant mediators alleviate the severity of DKD in mice^7–9, 16^. Thus, reducing inflammation may be a viable strategy to improve renal outcomes in patients with diabetes. However, there are no available kidney-enriched immune modulators.

Endothelial cell-specific molecule-1 (Esm-1), is a potential kidney-enriched immune modulator. Esm-1 is an endothelial-secreted circulating proteoglycan, notably expressed by the human glomerulus^17, 18^. Esm-1 was initially described as an inhibitor of the interaction between the leukocyte integrin LFA-1, and its endothelial ligand ICAM-1^19^, thus explaining its likely anti-inflammatory role in acute and chronic inflammatory conditions^20^. Esm-1 may attenuate inflammatory processes in DKD, as suggested by the higher *Esm1* expression levels observed in the glomeruli of mice resistant to albuminuria and DKD^18^. Our group has addressed the contribution of Esm-1 to the development of renal inflammation associated with DKD by using complementary gain- and loss-of-functional approaches. We found that in diabetic, Esm-1 deficiency mice, add back of Esm-1 induces protection from podocyte loss and albuminuria^21^. Several studies have also outlined a role for Esm-1 as a mediator of angiogenesis with a restricted expression to a minority of cells at the vascular tip^22–25^.

The precise anatomical and functional characteristics of *Esm1*(+) endothelial cells have not been described, and little is known about the role of this cell population in the mature kidney or in the development of DKD. We hypothesize that *Esm1* delineates a distinct subset of specialized cells in the glomerular vasculature, and that loss of *Esm1* expression in diabetes may also determine the transcriptional changes occurring in endothelial and neighboring glomerular cell types to mediate phenotypic changes in DKD.

In the present study, using single-cell RNA-sequencing, microarray data, and Esm-1 over-expression in an experimental model of DKD, we investigate the abundance and functional specialization of the *Esm1*(+) cell population in human and mouse mature kidney, we explore the influence of diabetes on its characteristics, and finally assess the effects of Esm-1 on the transcriptional and phenotypical features of DKD.

## Methods

### Bioinformatic analysis

#### Data collection and cluster identification

We downloaded gene raw counts or normalized gene expression matrix for each single-cell library from the supplemental material in Barry *et al*, 2019^26^, GEO databases (GEO Accession No. GSE129798; GEO Accession No. GSE131882; GEO Accession No. GSE140989), from the REC Dehydration database (https://endotheliomics.shinyapps.io/rec_dehydration/) or from the Human Kidney Cell Atlas (https://cellgeni.cog.sanger.ac.uk/BenKidney_v2.1/Mature_non_PT_parenchyma_v2.1.h5ad, mature human kidneys) (Table S1). For the glomerulus wide-genome expression profiling, we downloaded normalized processed data from GEO (GEO Accession No. GSE96804) (Table S2).

For the REC (Renal Endothelial Cell) Dehydration database (Database M1^27^), we downloaded normalized processed data filtered for high-quality cells from control mice. We then identified glomerular, cortical and medullary renal endothelial cells (gRECs, cRECs and mRECs) according to cluster annotations provided with the database. For the database from Barry *et al*, (Database M2^26^), we downloaded the raw counts of the scRNA-seq data, and following the criteria from the original publication, we analyzed cells that had between 200 and 2500 detected genes for downstream analyses to filter for high-quality single-cells^26^. We selected adult kidney cells according to cluster annotations provided with the database and normalized the expression matrix using NormalizeData with a scale set at 10,000 in the Seurat R package. We identified gRECs, cRECs and mRECs according to the expression of their top 50 marker genes on t-SNE projections using the Multigene expression tool in BD Genomics Dataview (version 1.2.2). For GSE129798 (Database M3^28^), we downloaded normalized processed data filtered for high-quality cells. We selected RECs according to the expression of the top 50 marker genes of RECs on t-SNE projections using the Multigene expression tool in BD Genomics Dataview. From the genesets we obtained the top 50 marker genes of RECs from the supplementary material in Ransick *et al*, 2019^28^. We identified cells from the Z2 and Z3 anatomical regions as mRECs. For RECs of the Z1 anatomical region, we identified gRECs and cRECs according to the expression of their top 50 marker genes on t-SNE projections using the Multigene expression tool in BD Genomics Dataview. We obtained the top 50 marker genes of gRECs and cRECs from the supplementary material in Dumas *et al*, 2020^27^.

For GSE131882, we downloaded index read count expression matrix from control (Database H1^29^) and diabetic subjects (Database H4^29^) and analyzed genes expressed in >3 nuclei and nuclei with at least 500 genes to filter data for high-quality cells, following the criteria from the original publication^29^. We did not retain data from Control 2 and Diabetes 2 subjects because of a high rate of nuclei not passing quality filtering. We then normalized the expression matrix using NormalizeData with a scale set at 10,000 in the Seurat R package (version 3.1.3) and restricted downstream analysis to RECs according to the expression of the canonical REC marker genes PECAM1 and FLT1 on t-SNE projections in BD Genomics Dataview. We then identified the various clusters of RECs (glomerular capillaries, afferent and efferent arterioles / peritubular capillaries, ascending and descending vasa recta and other ECs) according to the expression of the top 50 marker genes on t-SNE projections using the Multigene expression tool in BD Genomics Dataview. We labeled glomerular capillary cells as the gRECs subset, and every other EC as the cRECs subset. We obtained the top 50 marker genes of the REC clusters from the supplementary material in Lake *et al*, 2019^30^. For the Human Kidney Cell Atlas database (Database H2^31^), we downloaded normalized processed data filtered for high-quality cells. We selected RECs for downstream analysis and identified the various clusters (glomerular endothelium, peritubular capillary endothelium, ascending and descending vasa recta endothelium) according to the supplementary material in Stewart *et al*, 2019^31^. We labeled glomerular endothelium cells as gRECs, peritubular capillary endothelium as cRECs and ascending and descending vasa recta endothelium as mRECs. For GSE140989 (Database H3^32^), we downloaded normalized processed data filtered for high-quality cells. We selected RECs for downstream analysis and identified the various clusters (glomerular capillaries, peritubular capillaries, and arterioles afferent and efferent) according to cells annotations provided by the authors of the original publication^32^. We then normalized the expression matrix using NormalizeData with a scale set at 10,000 in the Seurat R package to adjust the normalization scale to that of other databases. Genesets of the top 50 marker genes used for the identification of each cluster in single-cell databases are described in Table S3^27, 30^. For GSE96804, we downloaded normalized processed data of glomerulus expression profiling on the Affymetrix microarray platform. For each subject, we obtained memberships to control (Database H5^33^) or diabetic (Database H6^33^) groups from the annotations provided with the database.

The main characteristics of each database are shown in **Tables S4** (single-cell) and **S5** (gene expression profiling).

#### t-SNE projections

We processed normalized data in BD Genomics Dataview to generate t-SNE projections. We used the *select variable genes* function to reduce the number of genes to a subset of most variable genes. We calculated the dispersion of log-transformed gene expression data for each gene and divided the genes into a number of bins based on mean gene expression per cell. In each bin, we calculated the z-score of the dispersion of each gene and retained genes with a z-score exceeding the dispersion threshold. We filtered out cells which had no molecule counts for the retained set of genes. We then calculated t-SNE coordinates on log-transformed data with a pseudo-count of 1 added using the *bh-t-SNE* function to construct a two-dimensional representation of the data (see **Table S6** for parameter settings for each database).

#### Single-cell expression analysis

We performed expression analyses on counts matrices scaled by total counts for each cell, multiplied by 10,000 and transformed to log space. We visualized the expression of *Esm1* and canonical genes of REC clusters on t-SNE-projections using the Gene expression in BD Genomics Dataview. We assessed the expression of canonical genes for various REC clusters in *Esm1*(+) cells through heatmap analysis. We produced heatmaps using the ggplot2 R package (version 3.3.0) on the basis of average log-transformed expression of the top 50 available marker genes (excluding Esm-1) according to previously published gene sets (see Table S3 for the detailed list of gene sets). We used the ggplot2 R package for visualization of log-transformed gene expression through violin plots.

#### Differential expression analysis

We used the *differential expression* function in BD Genomics Dataview to perform a negative binomial test on linear counts between *Esm1*(+) and *Esm1*(-) cells for all genes in each database.

#### Trajectory inference

We used the STREAM package in Python^34^ with default parameters to place cells onto pseudotime trajectories. We used a non-parametric local regression method (LOESS) to fit the relationship between mean and standard deviation values and selected the genes above the curve that diverged significantly as variable genes^35^. We performed dimensionality reduction using the Modified Locally Linear Embedding (MLLE) method^36^. We imported vasculature compartments as input data and subsequently visualized each cell membership onto linear pseudotime plots along with log-transformed expression of Esm-1. We subsequently compared scaled expression levels by Mann Whitney U tests.

#### Total RNA expression analysis

For the gene expression profile analysis of the human glomerulus (Database H6), we displayed the distributions of gene expression as boxplots using the ggplot2 R package and compared gene expressions between groups using the limma R package implementation in the Geo2R web tool.

For the gene expression profile analysis in mice with STZ-induced diabetes, we used the DEseq2 R package to process the data. We performed a shifted logarithm transformation of counts normalized on size factors to generate a log transformed count matrix for correlations analysis. We used the *rlog* function to generate a regularized log transformed matrix for Differential expression analysis.

#### Correlations analysis

We used the *corrplot* function from the corrplot R package to display correlation matrices between gene sets. When indicated, we performed hierarchical clustering of genes order using the hclust method to identify clusters of correlated genes. We used the ggplot2 R package to display heatmaps of correlations.

We compared the magnitude of the correlation patterns between the H5 and H6 databases by retaining the number of correlations with an arbitrarily fixed absolute r-value greater than or equal to 0.8 among all tested gene combinations.

For the untargeted correlation analyses in database H6, we retained genes with a q-value < 0.05 for differential expression in endothelial, mesangial and podocytes clusters from the supplementary material in Lake *et al*^30^ and in dendritic cells and macrophages from the supplementary material in Stewart *et al*^31^.

For the untargeted correlation analyses of the gene expression profiling in streptozotocin-induced DKD in mice, we retained genes with a q-value < 0.05 for differential expression in endothelial clusters from the supplementary material in Park *et al*^37^ and Ransick *et al*^28^, in podocytes and macrophages clusters from the supplementary material in Park *et al*^37^ and in mesangial clusters from Karaiskos *et al*^38^.

For selected genes of interest, we calculated and displayed linear regressions with 95% CI on scatter plots using the lm method in the ggplot2 R package. For analysis of the IGFBP5 interactome, we identified candidate genes using the Biogrid 3.5.185 website.

#### Gene and ontology enrichment analysis

We used the online software Metascape (http://metascape.org/) tool^39^ to perform single and multiple gene sets overlap analysis and identify enriched gene annotations across databases. We ran this analysis with default parameters to infer a hierarchical clusterization of significant ontology terms into a tree based on Kappa-statistical similarities among gene memberships. We selected the term with the best q-value within each cluster as its representative term. We used the 50 genes with top positive log fold-changes in counts between *Esm1*(+) and *Esm1*(-) cells and with a q-value of less than 0.05 as a multi-list input for Metascape enrichment analysis comparing databases H1, H2, H3, M1, M2, and M3. We used all differentially expressed genes in *Esm1*(+) cells with a q-value of less than 0.05 as single inputs for Metascape enrichment analysis comparing databases H1 and H4. For Metascape enrichment analysis of database H6, we used the genes respectively included in the negative and positive *Esm1* correlation clusters and with a q-value < 0.05 as a multi-list input. For Metascape analysis of the gene expression profiling of mice with STZ-induced DKD, we used the whole sets of genes belonging to the indicated clusters as single inputs (see Table S7 for gene lists used in Metascape analysis).

#### Gene lists overlap

We used the UpSetR Shiny web application (https://gehlenborglab.shinyapps.io/upsetr/)^40^ with standard parameters to visualize overlapping between lists of genes.

### Human samples

We obtained human biopsy specimens from neo-neoplastic tissue from nephrectomy specimens from a biorepository of patients with and without diabetes mellitus. We matched representative non-diabetic controls and diabetic samples for age and sex at the time of nephrectomy (Table S8).

### Animal models

#### Induction of diabetes and over-expression of circulating Esm1

We induced diabetes in eight week-old male DKD-susceptible (DBA/2) mice and used hydrodynamic tail-vein injection to over-express Esm1 as detailed elsewhere^41^. Data from this model obtained in diabetic mice without Esm1 over-expression (Database M4) and with Esm1 overexpression (Database M5) are accessible from Geobank with the accession number GSE175449.

#### Quantification of Albuminuria

We collected urine from mice in singly housed in metabolic cages (Tecniplast, Italy) as detailed elsewhere^21^.

#### RNA extraction for RNAseq analysis

We extracted kidney glomeruli from control, diabetic, and diabetic mice with Esm-1 over-expression for parallel analysis, as detailed elsewhere^41^. We isolated glomerular RNA by the Zymoresearch RNA preparation micro kit (Zymoresearch, Irvine, CA). We removed genomic DNA by RNase-free DNase I (Fermentas, Waltham, MA). We then reversed transcribed RNA into cDNA by using Improm II reverse transcriptase (Promega, Madison, WI) and random hexamer primers (Genelink, Elmsford, NY). We submitted cDNA samples to CD Genomics (New York, NY) for DNA quality control, library preparation, and sequencing (Illumina Novaseq PE150 sequencing, 20M read pairs).

#### In situ hybridization via RNAscope

We studied localization of target mRNA using the RNAscope Multiplex Fluorescent v2 kit (Advanced Cell Diagnostics, Newark, CA), according to the manufacturer’s instructions. We detected hybridization signals on 5-mm formalin-fixed paraffin-embedded transverse kidney sections using the TSA Plus fluorophores Opal 520 and Opal 690, with respective dilutions at 1:750 and 1:1500 (Akoya Biosciences, Waltham, MA). We mounted slices with ProLong Gold Antifade Mountant (Thermo Fisher Scientific, Waltham, MA) and viewed with a Leica DM5000 (Leica Microsystems, South San Francisco, CA). We included tissue from *Esm1* knockout mice^21^ and dapBcontrol probes as negative controls. Availability of probes limited our analysis. *Emcn, and Flt1* were the best glomerular capillary marker genes for whom an available probe could be used in mouse and human sections, respectively. Genesets of the top 50 marker genes used for the identification of cell subpopulations are described in Table S3^27, 30^.

### Study Approvals

The Stanford University School of Medicine Institutional Review Board (IRB) approved the collection and storage of human samples and clinical data from the Stanford Pathology Biobank, including a waiver of informed consent. The Stanford Research Compliance Office Administrative Panel for the Protection of Human Subjects – IRB approved the use of stored human biopsy samples. The Stanford Institutional Animal Care and Use Committee approved the experiments in mice.

### Statistical analysis

Details regarding the number of samples for each statistical comparison are provided in Table S4. For single-cell analysis, statistical significance was assessed with Chi2 square test using the *chisq.test* function in R when comparing categorical data. We performed binomial regressions on linear counts using the *differential expression* function in BD Genomics Dataview to compare single-cell gene expression levels. For the glomerulus gene expression profiling analysis, unpaired t-test were performed using the GEO2R web implementation of the limma R package with default settings. We run Spearman tests using the *corr.test* function from Psych R package to assess the correlation between two continuous variables. A bilateral p-value < 0.05 was considered statistically significant for all tests. When indicated, we calculated q-values using the *q-value* function from the q-value R package (version 2.18.0) with default parameters.

### Results

#### Esm1 is restricted to defined subsets of renal endothelial cells in mice and humans

To understand the features associated with the variation of *Esm1* expression in DKD, we first explored the characteristics of *Esm1* expression in healthy kidney vasculature. In mouse kidney, we first compared the expression of *Esm1* with canonical genes of vasculature compartments. *Esm1* is expressed in 12% to 18% of gRECs, 42% to 77% of cRECs, and 14% to 38% of mRECs (**Figure 1A** and Table S4). Subsequently, t-SNE projections show predominant expression of *Esm1* in regions enriched for *Igfp3, Igfbp5* or *Plpp3,* three markers of cRECs^27^ (**Figures S1A-C**). Congruent with this observation, *Esm1*(+) cells express higher levels of cRECs markers *Igfbp3*, *Igfbp5* and *Plpp3* (**Figure S1D**). Heatmap analysis of the top 50 marker genes of vasculature compartments confirms the preferential expression of cRECs markers (*Igfbp3, Igfbp5, AW112010, H2-K1, Rsad2, Kdr, Gpihbp1)* in *Esm1*(+) cells (**Figure 1B**). We performed localization of *Esm1*(+) cells by RNAscope. We observe preferential distribution of *Esm1*(+) cells in peritubular capillaries. We are also able to identify modest expression of *Esm1* in glomeruli (**Figure 1C**). However, further analysis reveals heterogeneity in transcriptomic signatures associated with the expression of *Esm1* within each compartment. Indeed, marker genes of afferent arterioles (*Sat1, Tmsb4x, Fabp4, Bst2, Ly6e, H2-K1)* and from the terminal portion of the afferent arterioles associated with the juxtaglomerular apparatus (*Tmem167b, Dll4, Cd300lg, Ifi44)* are co-expressed with *Esm1* in gRECs. In contrast, cells with glomerular capillary markers were not enriched for *Esm1* expression (**Figure S1E**). Furthermore, in cRECs, *Esm1* is mostly co-expressed with the top markers of cortical capillaries (*Ly6c1, Itm2b, Il10rb, Igfbp3, Igfbp5, Nrp1, Cyp26b1*), as well as markers of capillary angiogenic (*Gpihbp1, Fabp4, Sparc, Cd300lg*) and interferon markers (*Ifi44, Bst2, Rsad2*) (**Figure S1F**). Similar to cRECs, most capillary angiogenic (*Gpihbp1, Sparcl1, Fabp4, Sparc*) and interferon markers (*Rsad2, Ifi3m, B2m*) are enriched in *Esm1*(+) mRECs (**Figure S1G**).

**Figure 1.**
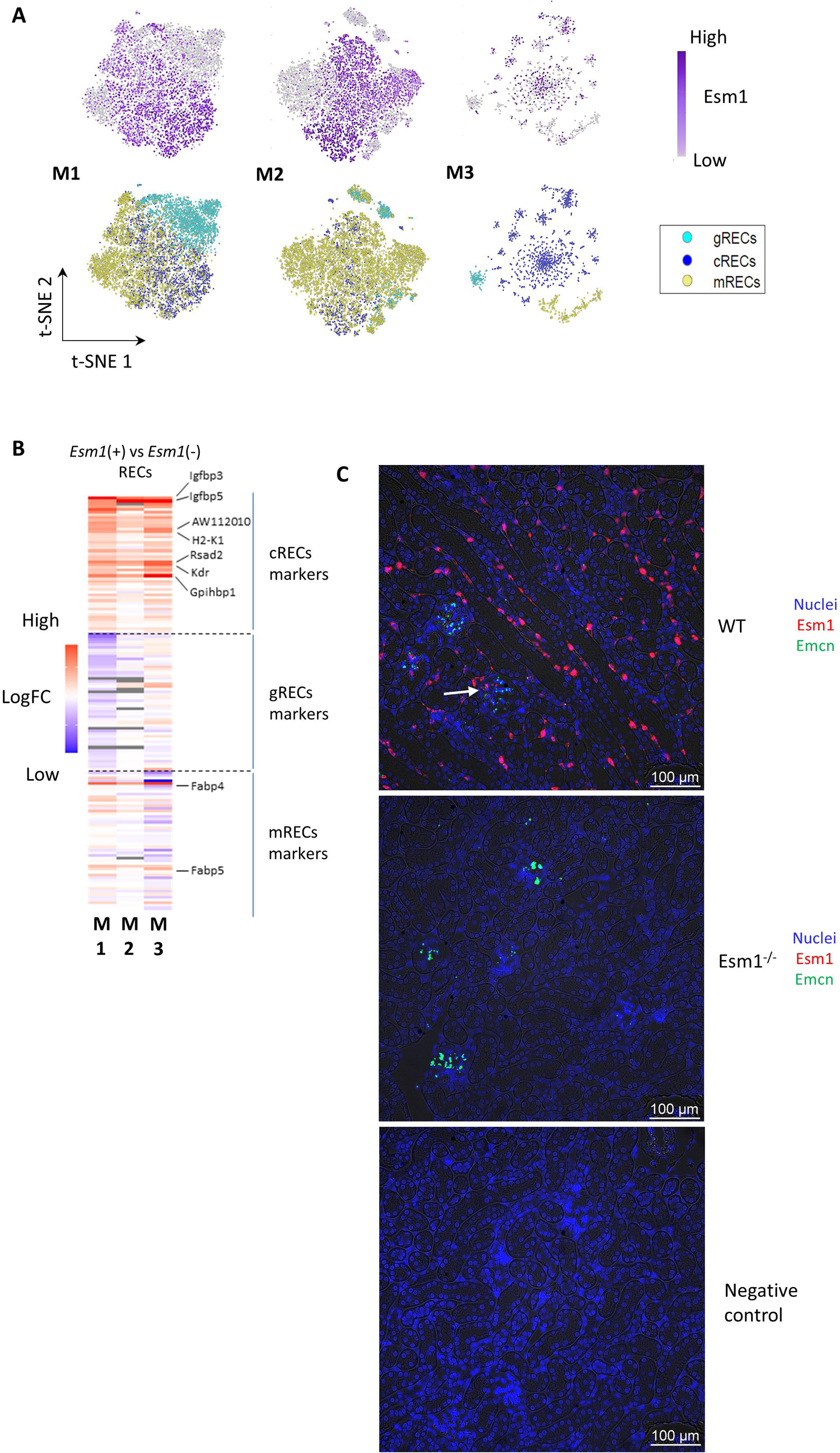
*Esm1* is expressed in various vasculature subsets including glomerular endothelial cells in mice. Data are from mice single-cell databases M1, M2, M3 for (A) and (B), and from C57BL/6 mouse kidney sections for (C). (A) t-SNE plots of RECs, are color-coded for the expression of *Esm1 (upper panel)* and for vasculature compartments *(lower panel)*. In the upper panel, no expression (*grey*), low (*light purple*) and high (*dark purple*) expression of *Esm1* are shown. In the lower panel, glomerular (g, *green*), cortical (c, *blue*), and medullary (m, *yellow*) are shown. (B) Heatmap of the relative fold-change in log-transformed expression-level of top 50 marker genes of mouse kidney vasculature compartments in *Esm1*(+) vs. *Esm1*(-) RECs. For heatmaps, blue is lower expression in *Esm1*(+) cells, red is higher expression in *Esm1*(+) cells, white is same expression in *Esm1*(+) and *Esm1*(-) cells. (C) RNAscope for *Esm1* (*red*) and *Emcn* (*green*) in the renal cortex. White arrows show glomerular expression of *Esm1*. REC, renal epithelial cells.

We then explored the characteristics of *Esm1*(+) cells in human kidney by comparing *Esm1* expression with that of canonical markers genes in vasculature compartments. *Esm1* is expressed in 23% to 51% of gRECs, vs. 13% to 33% of cRECs and 8% of mRECs (**Figure 2A** and Table S4). Accordingly, *Emcn* and *Syne1*, two glomerular capillaries markers, track with *Esm1* expression on t-SNE projections (**Figure S2A**) and are found at higher levels of expression in *Esm1*(+) cells in databases H2 and H3 (**Figure S2F**). Furthermore, we find enrichment of the top 50 glomerular capillaries markers (*EMCN, SYNE1, PTPRB, LDB2, FLT1, HSPA1A*) in *Esm1*(+) cells (**Figure 2B**). In human renal cortex, we observe that *Esm1* is mostly expressed in glomeruli, though modest expression is found in extra-glomerular cRECs (**Figure 2C**). These data show that glomeruli are the predominant source of renal *Esm1* expression in humans, in contrast to the mouse kidney where this compartment accounts for a minority of *Esm1* expression.

**Figure 2.**
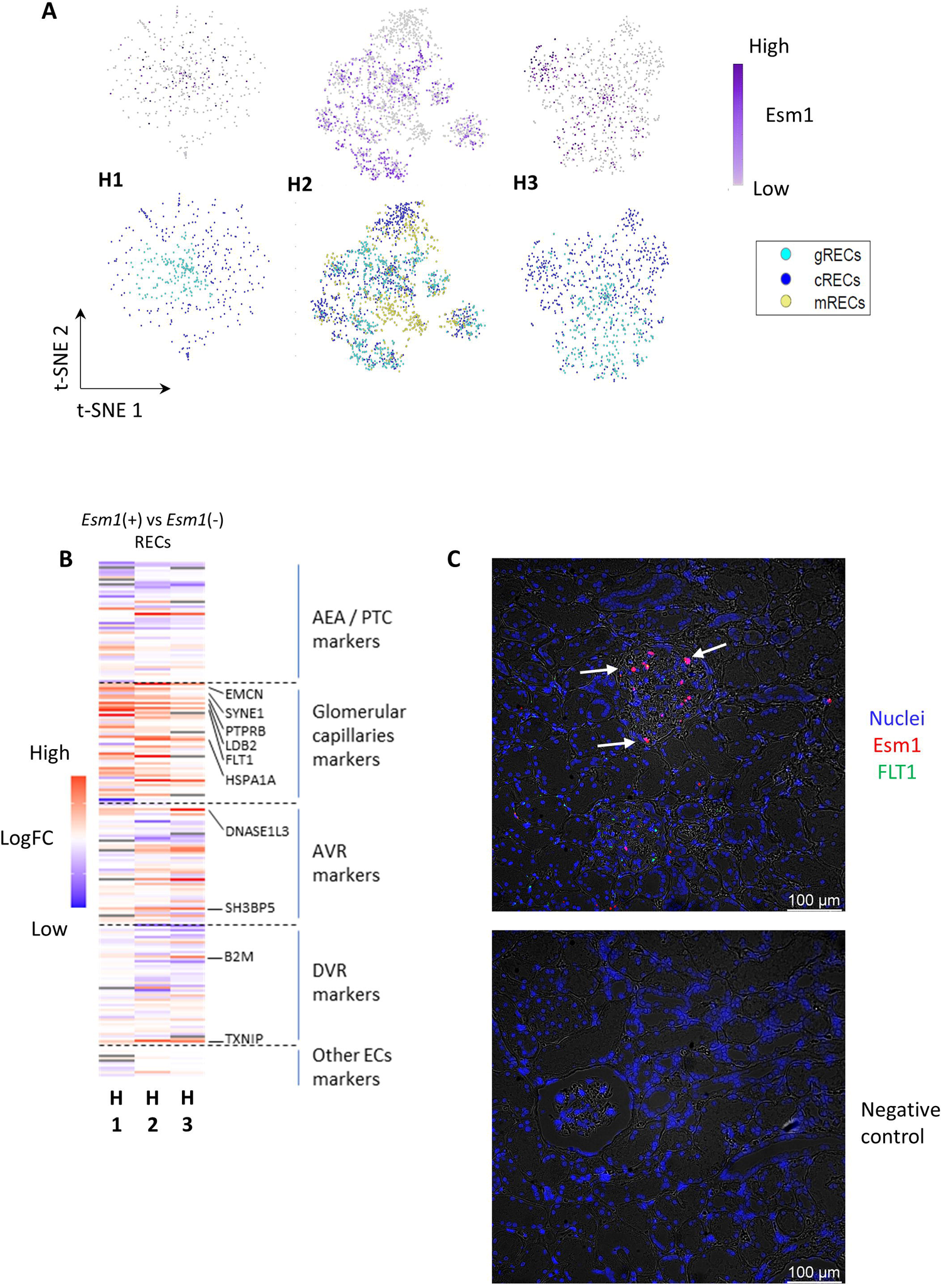
*Esm1* is mainly expressed in glomerular endothelial cells in humans. Data are from human single-cell databases H1, H2, H3 for (A) and (B), and from human kidney sections for (C). (A) t-SNE plots of RECs, are color-coded for the expression of *Esm1 (upper panel)* and for vasculature compartments *(lower panel)*. In the upper panel, no expression (*grey*), low (*light purple*) and high (*dark purple*) expression of *Esm1* are shown. In the lower panel, glomerular (g, *green*), cortical (c, *blue*), and medullary (m, *yellow*) are shown. (B) Heatmap of the relative fold-change in log-transformed expression-level of top 50 marker genes of human kidney vasculature compartments in *Esm1*(+) vs. *Esm1*(-) RECs. For heatmaps, blue is lower expression in *Esm1*(+) cells, red is higher expression in *Esm1*(+) cells, and white is similar expression in *Esm1*(+) and *Esm1*(-) cells. (C) RNAscope for *Esm1* (*red*) and *Flt1* (*green*) in the human renal cortex. White arrows show glomerular expression of *Esm1*. REC, renal epithelial cells.

#### Genes co-expressed with Esm1 differ across species, but are part of similar pathways

*Esm1* is not enriched in the same vascular compartments in mice and humans. Thus, we aimed to assess whether subsets of cells expressing *Esm1* are defined by common shared functions. Thus, we compared the top 50 marker genes in *Esm1*(+) (vs. *Esm1*(-)) cells in mouse and human kidney samples. We find that 13 of the *Esm1* co-expressed genes are shared between the three mouse gene lists, and 39 genes are found in common in at least two mouse gene lists. No co-expressed genes are shared between all three human gene lists; however, 12 genes are found in common in at least two human gene lists. Finally, 16 genes are shared across one mouse gene list and one human gene list. By contrast, gene ontologies are consistent across databases and species (**Figure 3A**). Four ontologies clusters, i.e., blood vessel development, epithelial cell proliferation, cellular response to growth factor stimulus, and morphogenesis of an epithelium, are significantly enriched in all mouse and human databases (detailed genes lists are shown in Table S7). Among these ontologies, blood vessel development is the most significantly enriched term, with 22% to 38% of the hits from each gene list falling into that term (enrichment score: 4.2 to 9.0; common q-value = 2.51*10^-12^) (**Figure 3B**). These four ontologies clusters are located in the same region on the gene ontology network, with a high similarity in ontology terms, as reflected by the number and thickness of edges linking each cluster to another (**Figure 3C**). We found several cluster nodes to be enriched for all human and mouse databases in each cluster node (**Figure 3D**) (see **Supplementary File 1** for interactive visualization of detailed enriched ontologies networks in Cytoscape). These results underline the biological processes shared by *Esm1*(+) cells, across multiple databases and species.

**Figure 3.**
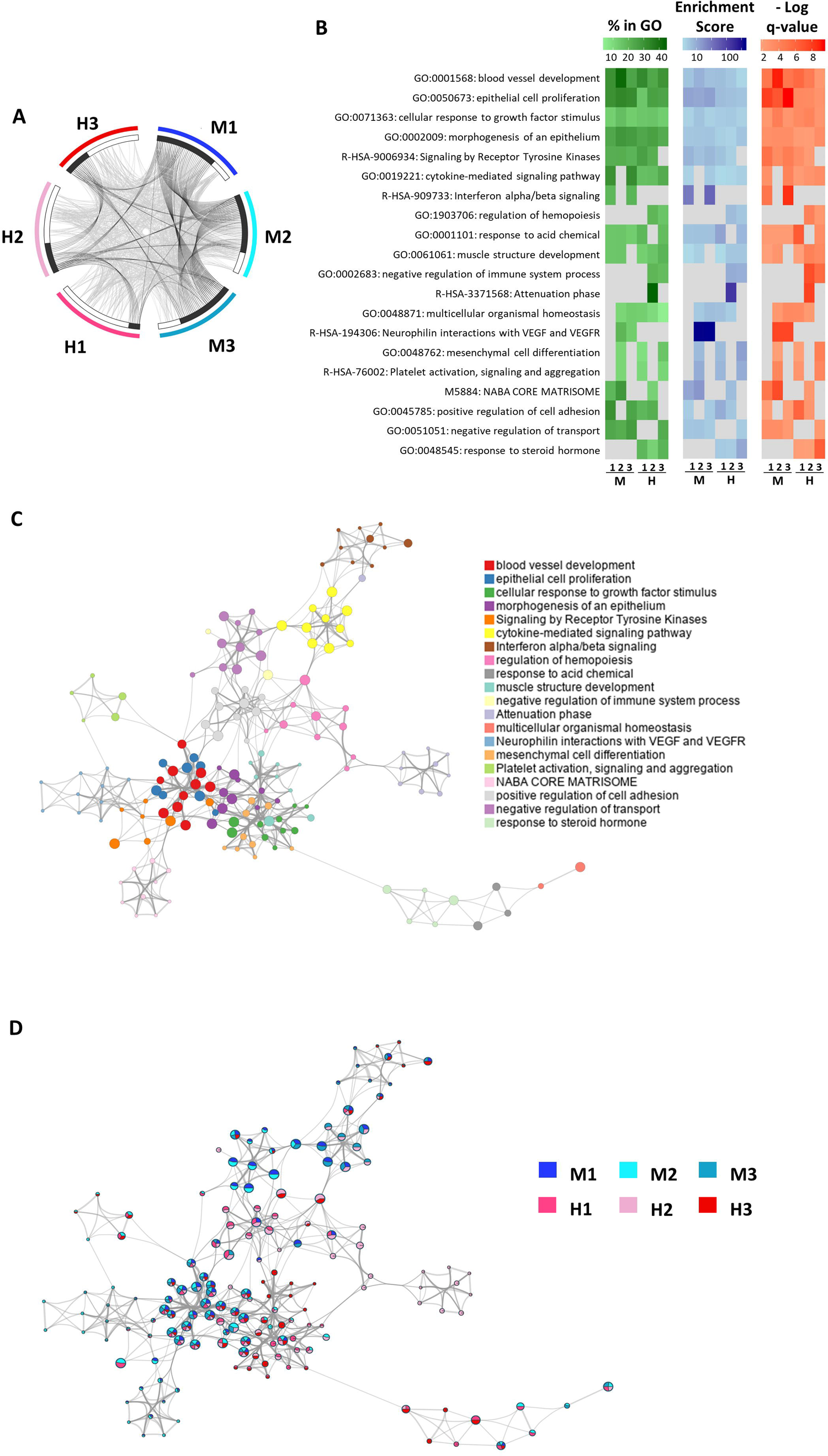
*Esm1*(+) cells share common functions across species. (A) Circos plot showing genes and annotations overlaps between gene lists expressed in Esm1(+) cells from each of the three mouse (M1-M3) and human (H1-H3) databases. Outer arcs represent the identity of each gene list. We represented each gene from a gene list as a spot on an inner arc. Genes that appear in multiple lists (*black*) and genes that are unique to the indicated list (*white*) are noted under each bar. Grey lines link the same gene that are shared by multiple gene lists. Black lines link the different genes falling into the same ontology term if statistically significantly enriched and with size no larger than 100. (B) Heatmaps of enrichment of ontology clusters, showing the hypergeometric q-values, enrichment scores and percentages of genes falling into each term for the top 20 most significantly enriched ontology cluster in each gene list. We performed a hierarchical clustering of significant ontology terms into a tree based on Kappa-statistical similarities among their gene memberships. We selected the term with the best q-value within each cluster as its representative term and display it in the heatmaps. Grey cells represent the non-significantly enriched ontology clusters in each gene list. The percentage of genes (*green*), enrichment score (*blue*), and q-value (*orange*) are shown for each ontology. (C) Network of enriched ontology terms. Each circle node represents a term, with the node’s size being proportional to the number of input genes falling into that term, and its color represents its ontology cluster identity, defined as the term with the best q-value within each cluster. Terms with a similarity score > 0.3 are linked by an edge, with the thickness of the edge proportional to the similarity score. The network is visualized with Cytoscape (v3.1.2) with “force-directed” layout and with edge bundled. (D) Network of enriched ontology terms with nodes displayed as pies. Each pie sector is proportional to the number of hits originated from a gene list. Color code for pie sector represents a gene list.

#### In diabetes, Esm1(+) cells undergo changes in relative Esm1 expression, abundance, and identity

To better understand the relationship between glomerular expression of *Esm1* and diabetes, we quantified the glomerular expression of *Esm1* in DKD-sensitive vs. resistant mice. Consistent with prior studies^18^, we observe lower glomerular intensity of *Esm1* in the DBA/2 DKD-sensitive, compared to the C57BL/6 DKD-resistant strain (average MFI: 0.09 vs. 0.14, p < 0.05) (**Figure S3A**). We also find lower intensity of *Esm1* from glomeruli of diabetic mice, compared to non-diabetic controls (average MFI: 0.03 vs. 0.09, p < 0.05) (**Figure S3B**). To verify these results in humans, we measured glomerular *Esm1* mRNA in 20 healthy controls and 41 patients with diabetes and find significantly lower expression with diabetes (log fold-change = −2.46, p = 4.06*10^-9^) (**Figure 4A**). In a separate cohort, we then quantified glomerular *Esm1* staining. Similar to results from mice, we observe reduced expression of human *Esm1* in diabetic patients, compared to healthy controls (average MFI: 0.01 vs. 0.21; p < 0.05) (**Figure 4B**).

**Figure 4.**
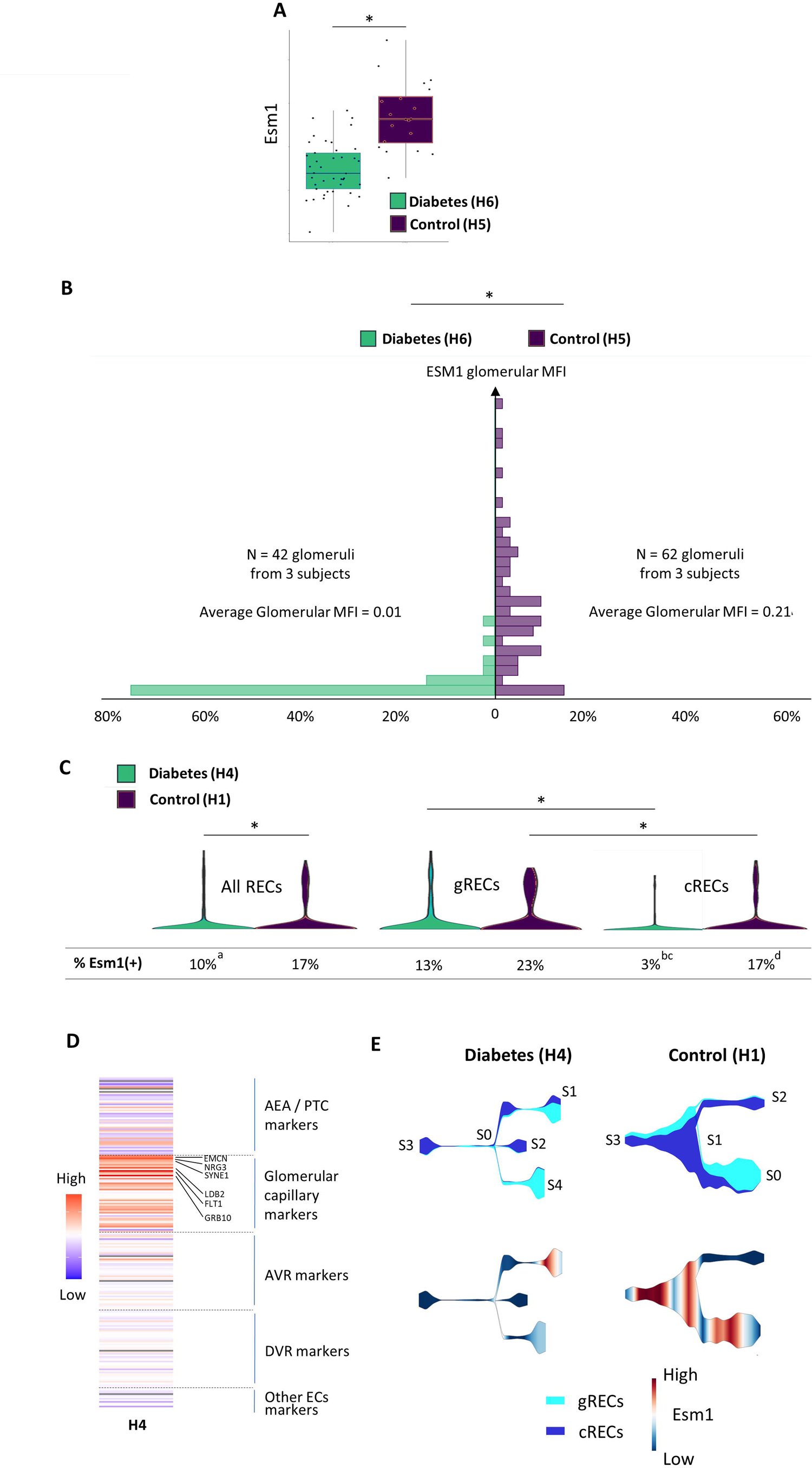
Glomerular expression of *Esm1* is decreased in diabetes and is associated with changes in specialization of *Esm1*(+) cells. (A) Total RNA glomerular expression of *Esm1* in 20 control and 41 diabetic subjects. Boxplots represent median [IQR]. (B) Quantification by RNAscope of human *Esm1* glomerular expression. (C) Violin plots showing the single-cell expression profiles of *Esm1* in human kidney vasculature compartments in control and diabetic subjects. Percentages of cells expressing *Esm1* are shown below violin plots. (D) Heatmap of the relative fold-change in log-transformed single-cell expression-level of top 50 marker genes of human kidney vasculature compartments in *Esm1*(+) vs. *Esm1*(-) RECs. (E) Visualization of inferred branching points, pseudotime trajectories, and expression of *Esm1* in diabetic and control subjects. We colored each zone according to cell cluster annotations (*upper panel*) or based on the expression of *Esm1* (*lower panel*). We annotated branching points and extremities arbitrarily. At a given pseudotime, the width of each branch is proportional to the total number of cells. In the lower panel, magnitude of *Esm1* expression is indicated by the scale provided. Data are from human genome expression profile databases H5 and H6 for (A), from 5µm human kidney sections for (B), and from human single-cell databases H1 and H4 for (C), (D) and (E). REC, renal epithelial cells. * p < 0.05 as indicated. ^a^ p < 0.05 vs. All RECs Control. ^b^ p < 0.05 vs. gRECs Diabetes. ^c^ p < 0.05 vs. cRECs Control. ^d^ p < 0.05 vs. gRECs Control.

To further characterize the transcriptional adaptation of human *Esm1*(+) cells to diabetes, we investigated the characteristics of *Esm1* single-cell expression in healthy controls and diabetic patients from two databases. Compared to cRECs, per-cell *Esm1* expression is higher and is detected in a greater percentage of cells in gRECs, both from controls (log fold-change = 0.09, p = 5.82*10^-4^; 23% vs. 13%, p = 0.005) and diabetic patients (log fold-change = 0.11, p = 2.98*10^-12^; 17% vs. 3%, p = 1.86*10^-6^) (**Figure 4C**). We find significant decreases in expression of *Esm1* (log fold-change = −0.079; p = 6.55*10^-25^) and % of *Esm1*(+) cells (10% vs. 17%; p = 0.004) from patients with diabetes, compared to healthy controls. Canonical markers of glomerular capillaries are also highly expressed in the *Esm1*-enriched region on t-SNE projections (**Figure S4A**), and in *Esm1*(+) cells (**Figure S4B**). Furthermore, *Esm1*(+) cells express most of the top 50 marker genes of glomerular capillaries (*EMCN, NRG3, SYNE1, LDB2, FLT1, GRB10*), by contrast with marker genes of other vasculature compartments (**Figure 4D**).

Additionally, we analyzed a pseudotime projection to infer the trajectory of cell specialization in the kidney vasculature. In diabetes, we observe most of *Esm1* expression at the extremities of the S0S1 and S0S4 diverging branches, delineating two subsets of specialized cells, mostly corresponding to glomerular capillaries (**Figure 4E**, **Figure S5A** and Table S9). In control subjects, *Esm1* is expressed in the S0S1 and S1S3 branches, two regions showing mixed enrichment for glomerular and cortical endothelial cells. Importantly, in contrast to the distribution observed in diabetes, *Esm1* is also expressed in the transition region connecting these two branches (**Figure 4E**, **Figure S5B** and Table S10). Taken together, these results demonstrate a reduction and restriction of *Esm1* expression to specialized subsets of glomerular endothelial cells in diabetes, compared with controls.

#### Diabsetes is also associated with a shift in the transcriptomic signature of Esm1(+) cells

To assess whether the lower abundance of *Esm1*(+) cells in diabetes is associated with a shift in function, we characterized the transcriptomic signature of these cells. *Esm1* expression is associated with a significant enrichment of 89 genes in healthy controls and 181 genes in patients with diabetes, with only seven genes overlapping between these two conditions (**Figure 5A**). While vascular development represents the ontology with the greatest number of *Esm1*(+) cell-enriched genes in controls, with 21 genes (enrichment score = 13; q-value = 1.4*10^-6^), chemotaxis has the most number of enriched genes in patients with diabetes, with 20 genes (enrichment score = 4.5, q-value = 1.26*10^-5^). The ‘positive regulation of vascular development’ ontology is not significantly enriched in *Esm1*(+) cells in diabetes (**Figure 5B**). Furthermore, we observe lower expression for most of the vascular development related genes in patients with diabetes, mostly driven by a lower expression in *Esm1*(+) than higher expression in *Esm1*(-) cells (**Figure 5C**). Conversely, in diabetes, we observe an increased enrichment in *Esm1*(+) cells for most genes in the chemotaxis ontology, due to higher expression in *Esm1*(+) cells and lower expression in *Esm1*(-) cells (**Figure 5D**). Taken together, these data underline a shift in the transcriptional program of *Esm1*(+) cells with diabetes.

**Figure 5.**
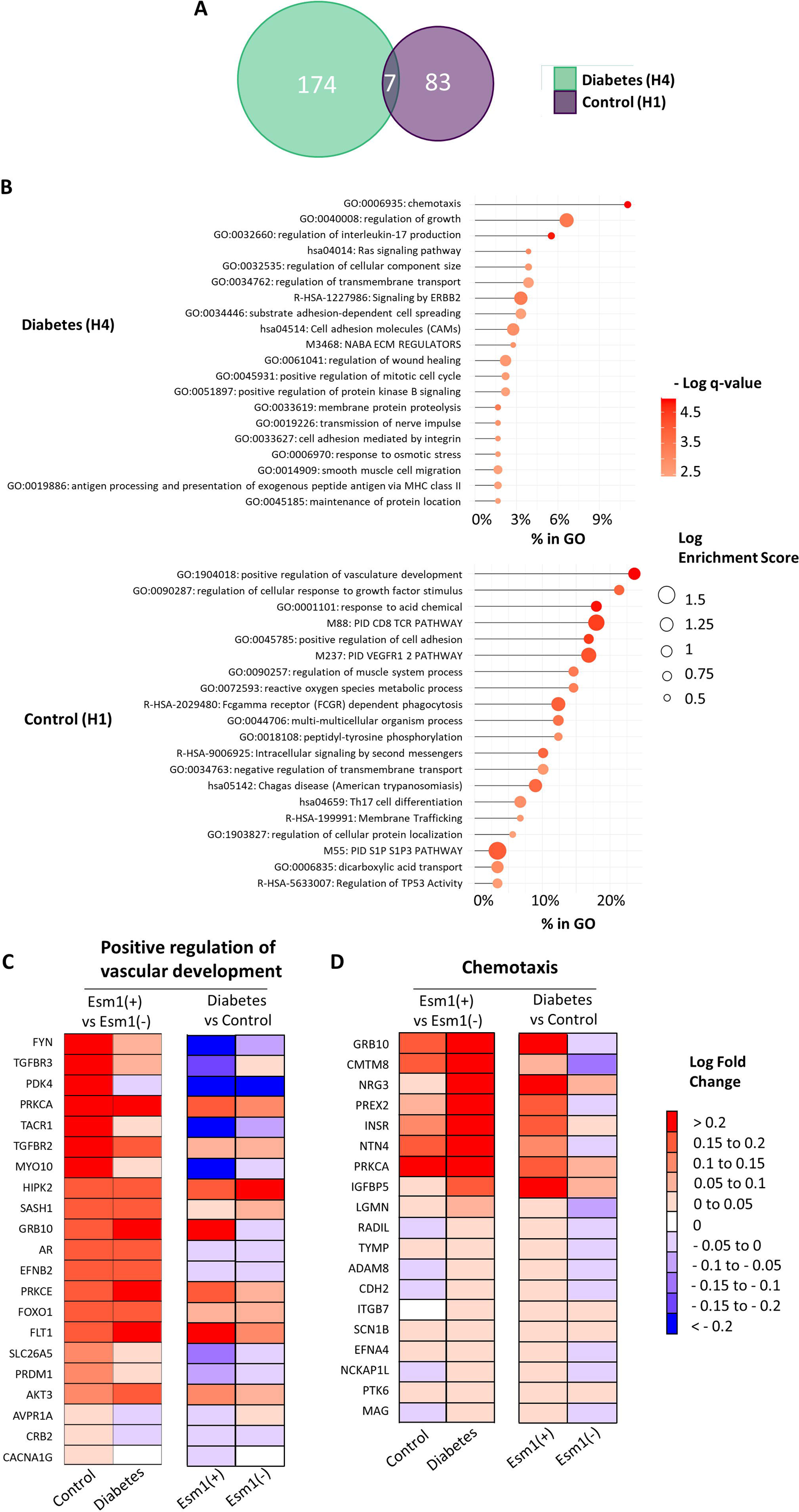
Diabetes is associated with a shift in the transcriptomic signature of *Esm1*(+) cells. (A) Venn diagram of the overexpressed genes in *Esm1*(+) cells in diabetic and control subjects. (B) Heatmaps of enrichment of ontology clusters, showing the hypergeometric q-values, enrichment scores and percentages of genes falling into each term for the top 20 most significantly enriched ontology cluster in each gene list. We performed a hierarchical clustering of significant ontology terms into a tree based on Kappa-statistical similarities among their gene memberships. We selected the term with the best q-value within each cluster as its representative term and display it on the graph. We ordered the ontology clusters by bar length representing the percentage of genes falling into each term. Enrichment scores are displayed as circle size and q-values as red gradient as indicated. (C) Heatmaps of the relative fold-change in *Esm1*(+) vs. *Esm1*(-) RECs (*left*) and in diabetes vs. control (*right*) in log-transformed expression-level of genes differentially expressed in *Esm1*(+) cells from control subjects falling into the Vascular development ontology cluster. (D) Heatmap of the relative fold-change in *Esm1*(+) vs. *Esm1*(-) RECs (*left*) and in diabetes vs. control (*right*) in log-transformed expression-level of genes differentially expressed in *Esm1*(+) cells from diabetic subjects falling into the chemotaxis ontology cluster. For heatmaps, relative expression in *Esm1*(+) vs. *Esm1*(-) cells is shown in controls (*left panel*) and diabetes (*right panel*) as indicated. All data are from human single-cell databases H1 (control) and H4 (diabetes). RECs, renal endothelial cells.

#### Esm1 single-cell co-transcriptome delineates glomerular tran in diabetes

We next assessed whether changes in the transcriptomic profile of *Esm1*(+) cells observed at the single-cell level could be confirmed in bulk transcriptomes. To that end, we analyzed a series of 20 healthy controls and 41 patients with diabetes for correlations between glomerular expression levels of *Esm1* and genes associated with the transcriptomic shift in *Esm1*(+) cells from our single-cell analysis. This analysis reveals that in diabetes compared with healthy controls, transcript levels within these gene clusters are significantly more correlated with each other and with *Esm1*. We observe a significant and direct correlation of glomerular *Esm1* with 14/21 vascular development genes. Accordingly, expression levels of these genes decrease along with that of *Esm1* in patients with diabetes. By contrast, these genes have a more stable glomerular expression level in healthy controls, regardless of *Esm1* expression, resulting in few correlations with *Esm1* (**Figures 6A** and S6A). In patients with diabetes, we observe a significant and direct correlation with *Esm1* for 7/8 genes that have a high single-cell co-expression with *Esm1*, which signifies that these chemotaxis genes have lower expression in diabetes, similar to *Esm1*. We also find a significant inverse correlation with *Esm1* for 6/11 genes with a low single-cell co-expression with *Esm1*, i.e., these genes have higher expression in diabetes, opposed to the decreased expression of *Esm1*. In contrast, in healthy controls, we find minimal correlation between expression of chemotaxis genes and *Esm1* (**Figures 6B** and **S6B**).

**Figure 6.**
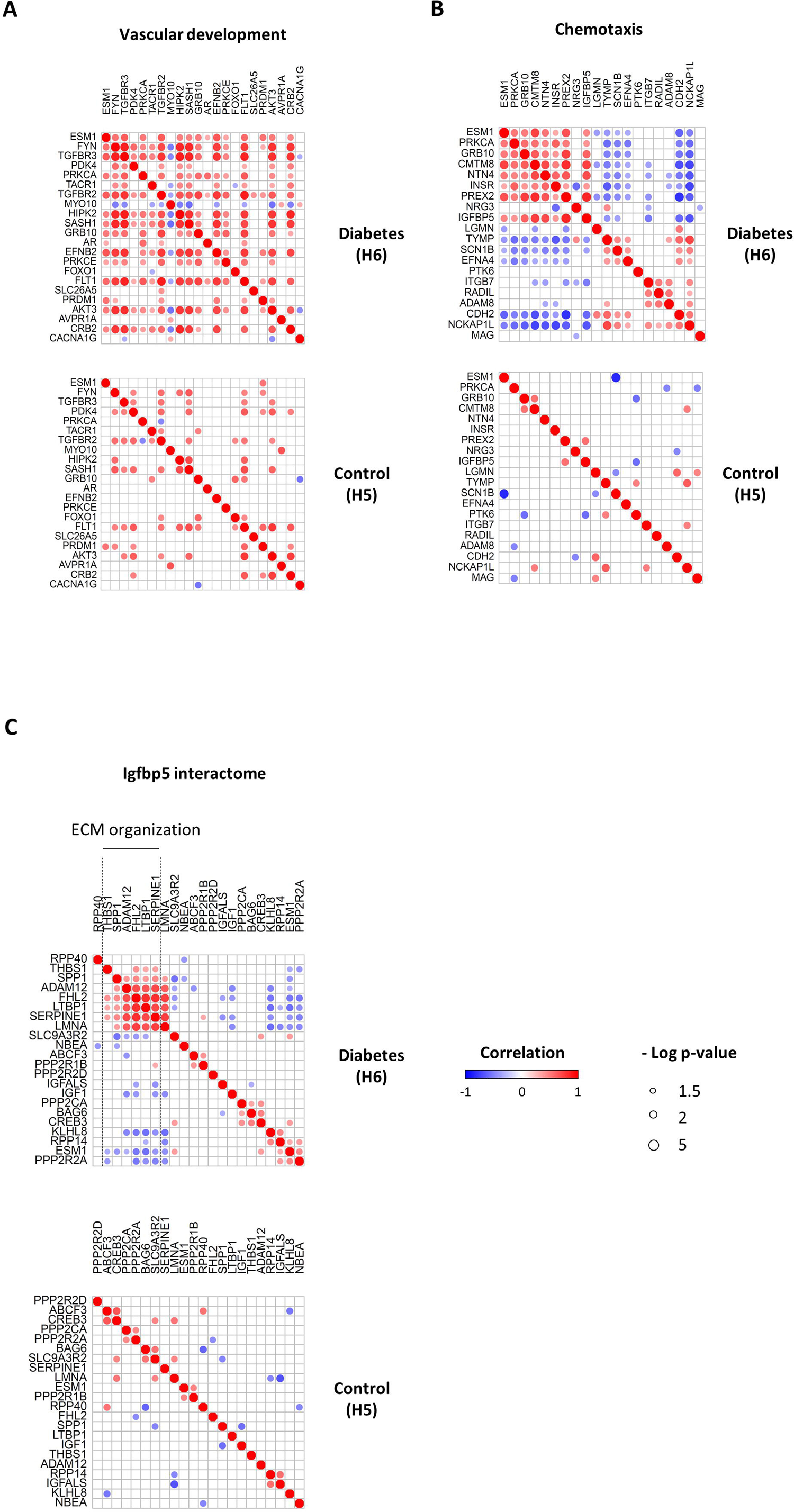
Human single cell diabetes-induced transcriptional shift in *Esm1*(+) cells observed in bulk transcriptomes. (A) Matrix of correlations of *Esm1* with genes differentially expressed in *Esm1*(+) cells from control subjects falling into the Vascular development ontology cluster. We ordered genes from higher to lower expression log fold-change in *Esm1*(+) vs. *Esm1*(-) cells in control subjects. (B) Matrix of correlations of *Esm1* with genes differentially expressed in *Esm1*(+) cells from diabetic subjects falling into the chemotaxis ontology cluster. We ordered genes from higher to lower expression log fold-change in *Esm1*(+) vs. *Esm1*(-) cells in diabetic subjects. (C) Matrix of correlations of *Esm1* with genes of the Igfbp5 interactome. We ordered genes following hierarchical clustering of correlations. For all correlation matrices, we displayed significant correlations as direct (*red circles*) or indirect (*blue circles*), with greater size representing more significant correlations. All data are from human gene expression profile databases H5 (control) and H6 (diabetes).

#### Potential cross-talk of Esm1(+) cells across glomeruli

To assess for cross-talk between *Esm1*(+) cells and other glomerular cell types, we examined gene expression among secreted genes. Of the chemotaxis genes that have higher expression in *Esm1*(+) cells from patients with diabetes, we further characterized *IGFBP5*. IGFBP-5 is an endothelial cell-expressed secreted protein, and we next analyzed for correlations between glomerular *Esm1* and the *IGFBP5* interactome in neighboring cells. We identified 21 genes coding for proteins with evidence of physical interaction with IGFBP-5. Among these, seven are found in a cluster of genes that all show a significant inverse correlation with *Esm1* in diabetes, i.e., the expression of these genes is higher in diabetic patients with lower *Esm1* glomerular expression. This cluster notably encompasses six genes from the extracellular matrix organization ontology, and 5/6 are abundantly expressed in neighboring mesangial cells (**Figures 6C** and **S6C**). Consistent with global changes in chemotaxis in diabetes, *Esm1*, through *IGFBP5*, links a subpopulation of endothelial cells with up-regulation of extracellular matrix genes enriched in mesangial cells.

#### Decreased Esm1 expression correlates with the level of transcriptional polarization in diabetes in individual glomerular cell types

To further study the relationship of *Esm1* with different glomerular cell types, we measured correlations of *Esm1* with genes enriched in specific cell types. For expression patterns of endothelial-enriched genes^30^, we find stronger correlation patterns in diabetes, with 253/22578 correlations identified with an absolute r-value ≥ 0.8, vs. 51/22578 correlations with an absolute r-value ≥ 0.8 in control subjects (p<0.05). We identify two clusters of genes in diabetes with direct and inverse correlations with *Esm1*, respectively, and these gene clusters are absent in healthy controls (**Figure 7A**). Among these clusters, we find a significant, direct correlation for 55 genes and a significant, inverse correlation for 36 genes from correlation clusters, respectively (Table S11). Blood vessel morphogenesis has the most number of genes directly correlated with *Esm1*, consistently with our previous findings of vascular development, encompassing 30/55 hits from this gene list (enrichment score = 11; q-value = 10^-13^). Nop56p-associated pre-rRNA complex is the only ontology cluster exclusively enriched with genes inversely correlated with *Esm1*, with 13/36 genes from this list (enrichment score = 45; q-value = 1.58*10^-10^). In addition, three ontology clusters are enriched for genes for both types of correlations, including RAGE signaling and extracellular matrix organization related pathways, suggesting that reduced *Esm1* expression in diabetes is associated with a more balanced effect on those pathways (**Figure 7B**). We then compared the expression levels of genes falling into the Nop56p-associated pre-rRNA complex and blood vessel morphogenesis between healthy controls and patients with diabetes and find that direct and inverse correlations with *Esm1* expression are observed in diabetes. Therefore, genes that inversely correlate with *Esm1* in diabetes exhibit either higher (e.g., *COL4A2*: log fold-change = 0.51, q-value = 1.08*10^-3^), similar (e.g., *IGFBP7:* log fold-change = 0.06, q-value = 0.4) or lower (e.g., *RPS16:* log fold-change = −0.29, q-value = 3.36*10^-6^) expression compared to the control group. Similarly, genes that directly correlate with *Esm1* in diabetes show either higher (e.g., *DOCK4:* log fold-change = 0.35, q-value = 1.27*10^-2^), similar (e.g., *PIP4K2A:* log fold-change = −0.15, q-value = 0.32) or lower (e.g., *TGFBR2:* log fold-change = −0.96, q-value = 4.77*10^-5^) expression compared to the control group (**Figure 7C**).

**Figure 7.**
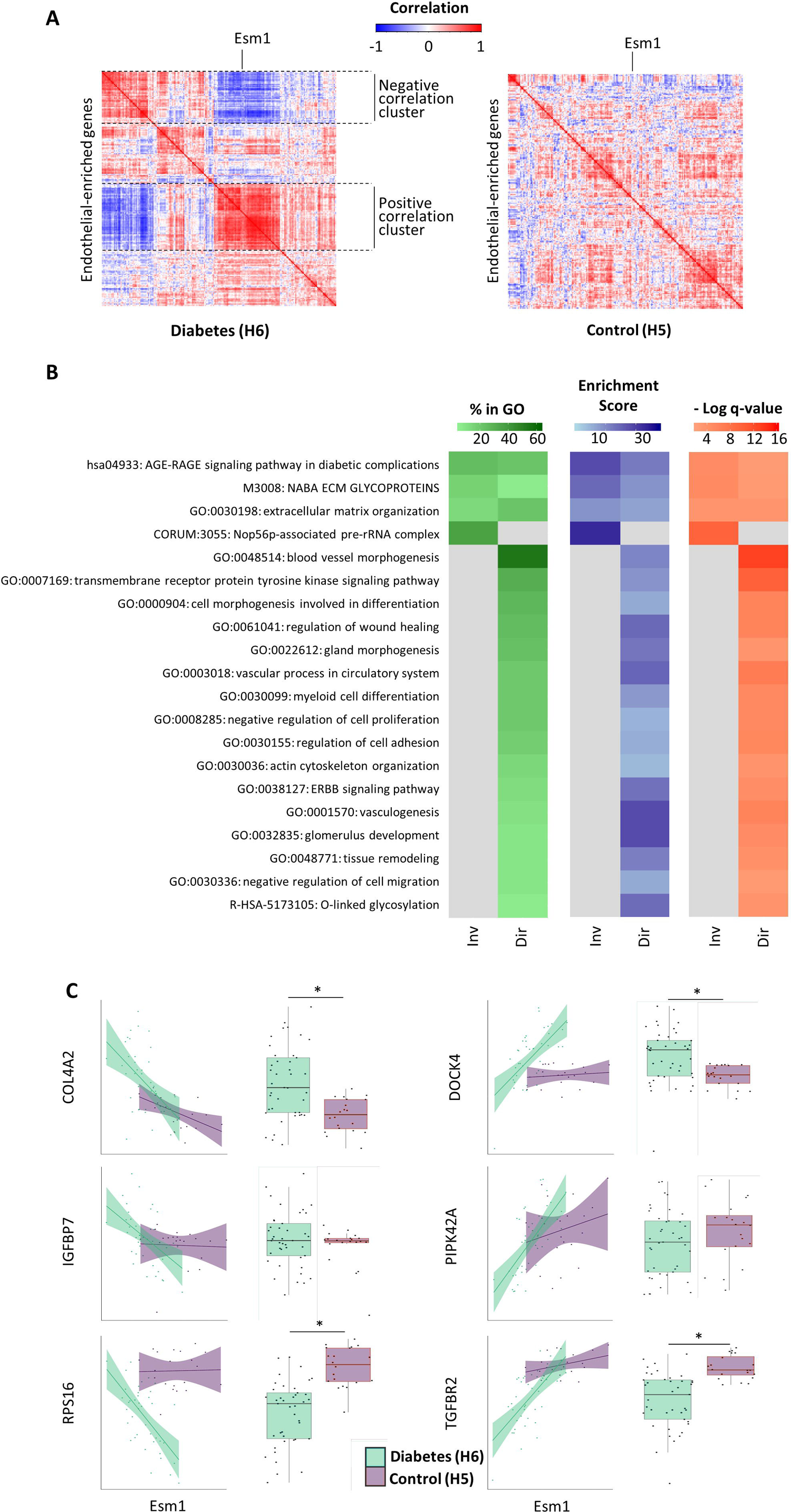
*Esm1* depletion is a sensor of the transcriptomic polarization of endothelial-enriched genes in diabetes. (A) Matrix of correlations of endothelial-enriched genes in healthy controls and patients with diabetes. We ordered genes by hierarchical clustering of correlations based on the color key as indicated. (B) Heatmaps of enrichment of ontology clusters, showing the hypergeometric q-values, enrichment scores and percentages of genes falling into each term for the top 20 most significantly enriched ontology cluster in each gene list among endothelial-enriched genes in patients with diabetes. We performed a hierarchical clustering of significant ontology terms into a tree based on Kappa-statistical similarities among their gene memberships. We selected the term with the best q-value within each cluster as its representative term and display it in the heatmaps. Grey cells represent the non-significantly enriched ontology clusters in each gene list. The percentage of genes (*green*), enrichment score (*blue*), and q-value (*orange*) are shown for each ontology. (C) Expression levels in control and diabetes of genes correlated to *Esm1* in diabetes, displayed vs. *Esm1* (*scatter plots, left panels*) and alone (*boxplots, right panels*). For each group, linear regression curve with 95%CI and median [IQR] are displayed on scatter plots and boxplots, respectively. Inverse (Inv) and direct (Dir) correlations with *Esm1* expression in diabetes are shown. All data are from human gene expression profile databases H5 (control) and H6 (diabetes). * p < 0.05 as indicated.

We next studied expression patterns of genes enriched in neighboring cell types, i.e., mesangial cells^30^, podocytes^30^, and immune cells^31^. In mesangial-enriched genes, we find stronger correlation patterns in diabetes, with 105/9591 correlations identified with an absolute r-value ≥ 0.8, vs. 27/9591 correlations with an absolute r-value ≥ 0.8 in control subjects (p<0.05). Further, we find two clusters of genes in diabetes with direct and inverse correlations with *Esm1* (**Figure 8A**). Among these, there is a significant correlation for 31 and 16 genes of the direct and inverse correlation clusters, respectively (Table S11).

**Figure 8.**
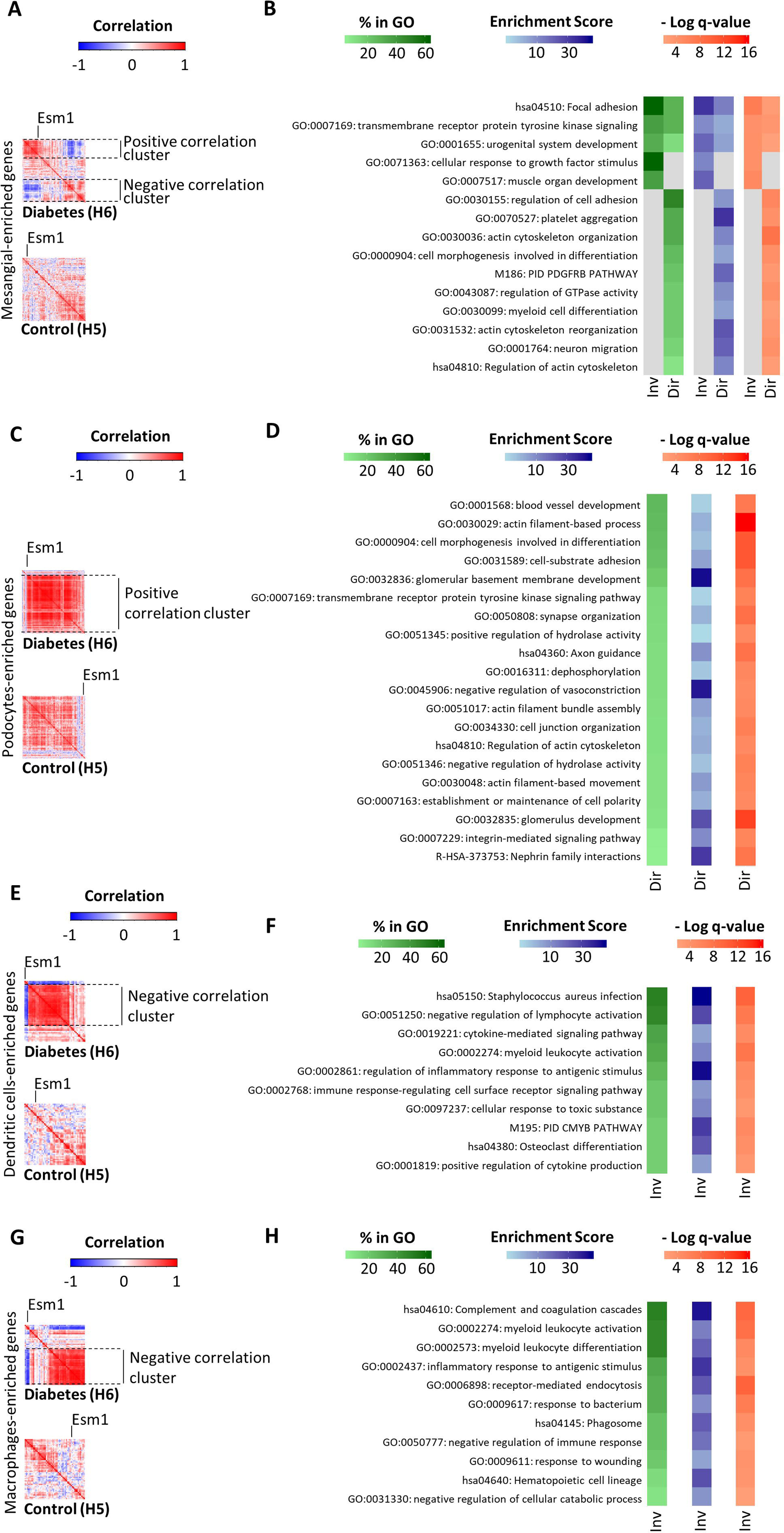
*Esm1* depletion is a sensor of the transcriptomic polarization of neighboring glomerular cell type specific-enriched genes in diabetes. Matrix of correlations of glomerular cell type specific-enriched genes in healthy controls and patients with diabetes (A, C, E, G) and heatmaps of enrichment of ontology clusters for select ontology clusters significantly enriched in at least one gene list of correlation among cell type-specific enriched genes in patients with diabetes (B, D, F, H). Mesangial cells (A, B), podocytes (C, D), dendritic cells (E, F), and macrophages (G, H) are shown. For all matrices of correlations, genes are ordered following hierarchical clustering of correlations. Red is for direct correlations, blue is for inverse correlations, and white indicates no correlation. For all heatmaps we show the hypergeometric q-values, enrichment scores and percentages of genes falling into each term. We performed a hierarchical clustering of significant ontology terms into a tree based on Kappa-statistical similarities among their gene memberships. We selected the term with the best q-value within each cluster as its representative term and display it in the heatmaps. Grey cells represent the non-significantly enriched ontology clusters in each gene list. The percentage of genes (*green*), enrichment score (*blue*), and q-value (*orange*) are shown for each ontology. Inverse (Inv) and direct (Dir) correlations with *Esm1* expression in diabetes are shown. All data are from human gene expression profile databases H5 (control) and H6 (diabetes).

Regulation of cell adhesion is the ontology cluster with the most number of genes directly correlated with *Esm1* (15/31 genes; enrichment score = 8.7; q-value = 2.51*10^-6^), while cellular response to growth factor stimulus encompasses the most number of genes inversely correlated with *Esm1* (10/16 genes; enrichment score = 11; q-value = 6.31*10^-5^). Furthermore, we find three ontology clusters enriched for genes from both lists (focal adhesion, transmembrane receptor protein tyrosine kinase signaling and urogenital system development) (**Figure 8B**). In podocytes-enriched genes, we find stronger correlation patterns in diabetes, with 5166/21736 correlations identified with an absolute r-value ≥ 0.8, vs. 269/21736 correlations with an absolute r-value ≥ 0.8 in control subjects (p<0.05). In diabetes, we only identify one cluster of genes in diabetes with direct correlations with *Esm1* (**Figure 8C**), among which 180 are statistically significant (Table S11). Blood vessel development is the ontology cluster with the most number of genes directly correlated with *Esm1* (48/180 genes; enrichment score = 3.8; q-value = 7.94*10^-8^). (**Figure 8D**). In dendritic cells and macrophages-enriched genes, we find stronger correlation patterns in diabetes, with 198/1225 and 218/1275 correlations identified with an absolute r-value ≥ 0.8, vs. 7/1225 and 6/1225 correlations with an absolute r-value ≥ 0.8 in control subjects (p<0.05), respectively. Dendritic cells and macrophages-enriched genes each have a cluster of indirect correlation with *Esm1* in diabetes (**Figures 8E** and **8G**), including 34 and 30 significant correlations, respectively (Table S11). *Staphylococcus aureus* infection has the most number of hits from dendritic cell-enriched genes (17/34 genes, enrichment score = 76; q-value 1.58*10^-10^) (**Figures 8F**) while complement and coagulation cascades encompass the greatest number of hits from macrophage-enriched genes (15/30 genes; enrichment score = 61, q-value = 10.31*10^-10^) (**Figures 8H**, intersections between lists of genes enriched in human glomerular compartments are shown in Figure S7). Taken together, across glomerular cell type-specific genes, we observe a consistent polarization in the transcriptomic profile in diabetes, and that these clusters correlate closely with changes in *Esm1* expression.

#### Over-expression of Esm-1 modulates the glomerular transcriptional shift associated with albuminuria in DKD

To understand the significance of glomerular transcriptional modifications in DKD, we studied the expression profile of genes enriched in glomerular compartments in a murine model of DKD with reduction in albuminuria (Database M4) and with over-expression of *Esm1* (Database M5)^41^. In 234 of 300 endothelial-enriched genes^28, 37^, we observe correlations between *Esm1* expression and albuminuria (κ = 0.53; p = 2.29*10^-29^) (**Figure 9A** and Table S12), i.e., genes that are directly correlated with *Esm1* show an inverse correlation with albuminuria, and those showing an inverse correlation with *Esm1* directly correlate with albuminuria. Furthermore, for 206 of these 234 correlation genes, systemic over-expression of Esm-1 reverses the expression profile (κ = 0.25; p = 3.4*10^-6^) (**Figure 9A** and Table S12). Many of the genes that directly correlate with albuminuria and are down-regulated by Esm-1, are from a vasculature development ontology (**Figure 9A**, Tables S7 and **S12**). Conversely, we find another cluster of genes that inversely correlate with albuminuria and are up-regulated by Esm-1 are related to blood vessel morphogenesis (**Figure 9A**, Tables S7 and **S12**). We find similar patterns of genes with an inverse correlation between glomerular *Esm1* and albuminuria that are enriched in different glomerular cell types: mesangial cells (35/49 genes; κ = 0.5; p = 4.1*10^-7^), podocytes (210/271 genes; κ = 0.49; p = 3.33*10^-23^) and macrophages (55/81 genes; κ = 0.4; p = 7.7*10^-6^) (**Figures 9B-D** and Table S12). Moreover, a large subset of these genes are directly regulated by Esm-1: mesangial (33/49 genes; κ = 0.37; p = 7.7*10^-4^) and podocytes-enriched genes (188/271 genes; κ = 0.21; p = 1.3*10^-4^) but not in macrophage-enriched genes (38/81 genes; κ = 0.04; p = 0.64) (**Figures 9B-D** and Table S12). Subsequently, we identify clusters of genes with inverse correlations with *Esm1*, direct correlations with albuminuria and that are downregulated with Esm-1 over-expression, mainly enriched for the regulation of vascular smooth muscle cell proliferation in mesangial-enriched genes, for actin filament-based process and ECM organization in podocytes enriched-genes and for the *Staphylococcus aureus* infection pathway in macrophages-enriched genes (**Figure 9B-D**, Tables S7 and **S12**). On the other hand, we find clusters of genes with direct correlations with *Esm1*, inverse correlations with albuminuria and that are upregulated with Esm-1 over-expression, mainly enriched for the regulation of vascular smooth muscle contraction in mesangial-enriched genes, for the regulation of cellular response to growth factor stimulus in podocytes enriched-genes and for myeloid leukocyte differentiation in macrophages-enriched genes (**Figures 9B-D**, Tables S7 and **S12,** intersections between lists of genes enriched in mouse glomerular compartments are shown in Figure S9). Taken together, over-expression experiments with Esm-1 demonstrate that for a large subset of glomerular cell type-specific genes that relate to albuminuria, Esm-1 determines expression of these genes.

**Figure 9.**
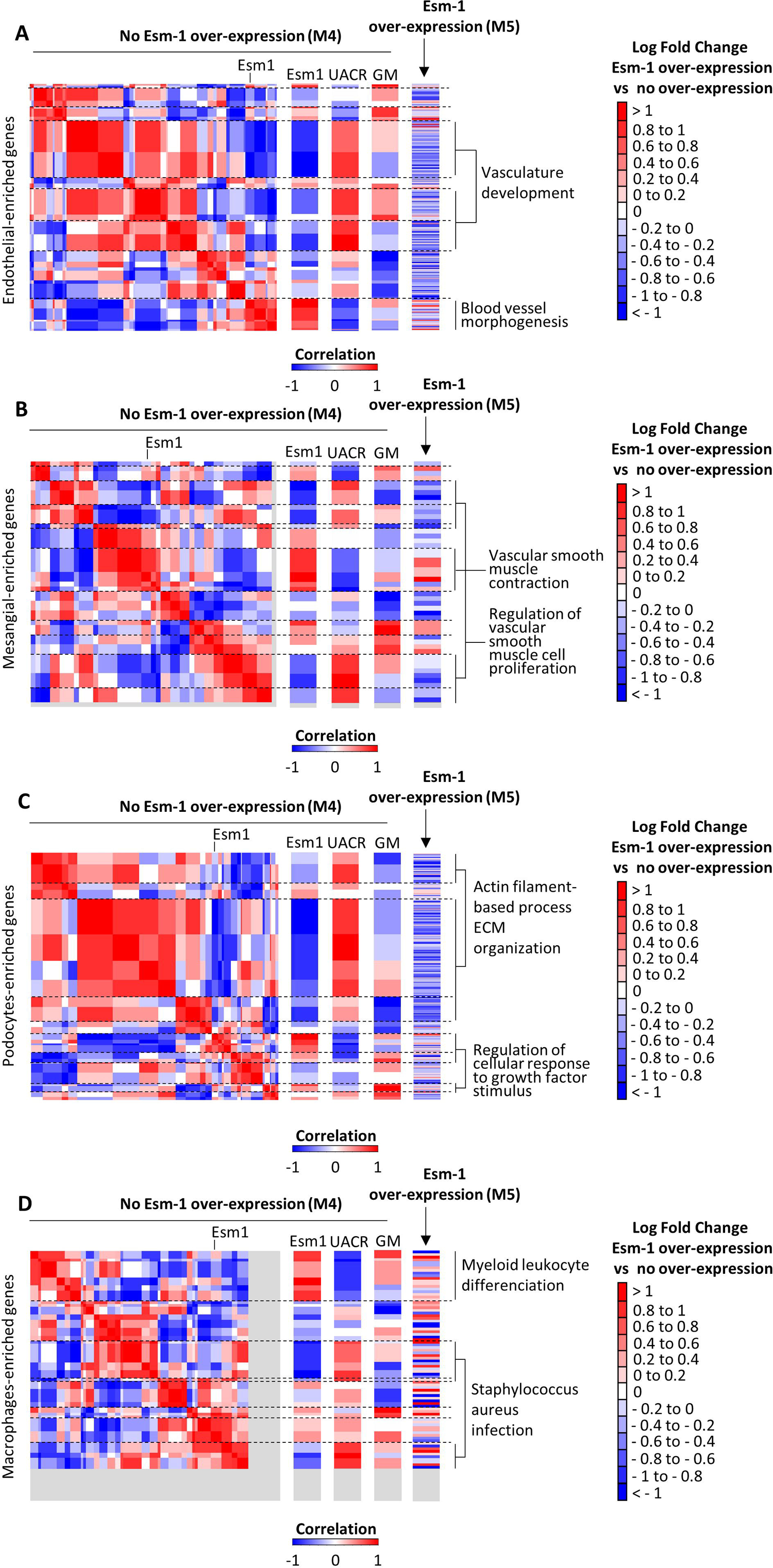
Systemic overexpression of mouse Esm-1 regulates the expression of genes correlating with albuminuria in mice with DKD. Matrix of correlations of glomerular cell type-specific-enriched genes, heatmaps of correlations with *Esm1*, urine albumin-to-creatinine ratio (UACR), glomerular macrophages (GM) and heatmap of expression log fold-change in mice with mouse Esm-1 systemic overexpression vs. no overexpression. Data for endothelial (A), mesangial (B), podocytes, and (C), and macrophages (D) are shown as indicated. For correlations matrices and heatmaps, red is for direct correlations, blue is for inverse correlations, and white indicates no correlation. For expression log fold-change heatmaps, blue is lower expression in mouse Esm-1 overexpression, red is higher expression in mouse Esm-1 overexpression, and white indicates no significant differences in expression with over-expression of mouse Esm-1. N = 4 in mice with DKD and no overexpression of mouse Esm-1. N = 3 in mice with DKD and overexpression of mouse Esm-1. All data are from DBA/2 mice with streptozotocin-induced DKD (Databases M4 and M5, see Zheng, et al., for details,^41^).

## Discussion

We show that *Esm1*(+) cells segregate as a distinct subset in each vasculature compartment, thus defining a specialized population of cells. Combined with the variety of the vascular compartments enriched for *Esm1*, intra- and inter-species, our data suggest that the transcriptomic signature of *Esm1*(+) cells may not be solely explained by their spatial localization. Our results highlight a common set of functions in mature kidney shared by this subpopulation, including but not limited to angiogenesis, thus designating *Esm1*(+) cells as playing a role in the tissue proliferation process. These results are consistent with previous observations showing the critical role of *Esm1* in vasculogenesis, with high expression at the vascular tip^21–24^. A common set of functions based on distinct co-expressed genes has been described before^42^, but has not previously been associated with *Esm1*, endothelial cell subtypes, or kidney-specific genes. *Esm1*(+) renal endothelial cells from patients with diabetes exhibit lower expression of a majority of genes involved in vasculature development. Taken together, these results may reflect an impaired ability to regenerate injured renal endothelium in diabetes due to various potential causes including: (1) a decrease in the pool of cells specialized in the tissue and endothelial repair process; and/or (2) a loss of function of this particular cellular subset. However, further studies are needed to assess if these impaired pathways are a cause or a consequence of fewer *Esm1*(+) cells.

On the other hand, we observe a distinct shift in the role of *Esm1*(+) cells from patients with diabetes. This shift, characterized by upregulation of chemotaxis-related genes, despite occurring in a minority population of endothelial cells, is reflected in the glomerular transcriptome by a lower expression of chemotaxis-related genes that are highly correlated with *Esm1* expression in subjects with lower expression levels of *Esm1*. Esm-1 transcription is up-regulated by several pro-inflammatory mediators^17^. Thus, one may propose that *Esm1* expression persists in a subset of cells involved in the inflammatory process. Although their cell numbers are lower with diabetes, higher expression of chemotaxis-related genes in *Esm1*(+) cells compared to other glomerular endothelial cells, may indicate a role for these cells to provide a homing signal for endothelial progenitor cells to attenuate glomerulosclerosis and renal fibrosis^43^. In addition, this shift in the pattern of *Esm1*(+) cell transcription can be linked to the inhibitory role of *Esm1* in the leukocyte recruitment process^18, 19, 44, 45^ in specific disease states. Therefore, the higher expression of *Esm1* in cells involved in the chemotactic process may reflect a potential mechanism for regulating vascular inflammation, thus supporting the hypothesis of its renoprotective effect in the diabetic kidney.

More generally, our results highlight the transcriptional polarization which occurs in the glomerular vasculature in diabetes. We observe organized alignments in the variations of gene expression that are not found in healthy controls. Interestingly, despite a loss of *Esm1*(+) cells in diabetes and lower expression of *Esm1*, *Esm1* appears to be a marker of this transcriptional polarization, as most variations in gene expression indirectly correlate with the intensity of glomerular *Esm1*. This transcriptional polarization is observed not only in glomerular endothelial cells, but also in neighboring glomerular cell types, thus raising the possibility of cross-talk between *Esm1*(+) cells and other glomerular cell populations in DKD. This finding is consistent with other studies showing evidence of intra-glomerular cell cross-talk^29, 46–49^. Given the ontologies aligning with *Esm1* expression in our study, we speculate that this communication network could provide feedback that may enable cells to tune their signaling activity to regulate specialized functions, including vascular growth and chemotaxis. Future studies on glomerular cross-talk will explore autocrine and paracrine effects of secreted factors by *Esm1*(+) cells, including IGFBP-5.

Interestingly, our study highlights a correlation between *Esm1* and *IGFBP5* downregulation in diabetes. Given that IGFBP-5 is secreted, we investigated possible associations with the transcriptomic profile of the *IGFBP5* interactome in neighboring glomerular cells. Based on this analysis, we find in patients with DKD with lower *Esm1* expression a consistent upregulation of *THBS1, SPP1, ADAM12, FHL2, LTBP1* and *SERPINE1*^50–53^, which are all involved in the regulation of *IGFBP5* in the extracellular matrix. Except for *SPP1*, all these genes have an enriched mesangial expression^29^, thus highlighting that variations in the *Esm1(+)* cell transcriptome might affect gene expression in neighboring glomerular cells. Interestingly, when analyzing the expression of endothelial genes correlating with *Esm1* expression, extracellular matrix pathways are found among the top enriched ontologies.

Another novel finding from our study is the relationship between the glomerular transcriptomic polarization and albuminuria. To the best of our knowledge, our study is the first to identify a systematic orientation of the glomerular-enriched transcriptome in which expression of genes align in parallel with the severity of albuminuria and with correlation to *Esm1* expression in DKD. We therefore determined next whether *Esm1* is a marker or mediator of this polarization. We identify subsets of these genes and pathways correlated to albuminuria in which gene expression is reversed by systemic over-expression of Esm-1. Importantly, we find that Esm-1 attenuates transcription of podocyte-enriched genes that are related to the extracellular matrix and are associated with high levels of albuminuria. This latter result is consistent with published data showing the protective effect of Esm-1 against podocyte loss^41^ and provides novel insights regarding potential molecular targets to prevent the development of DKD. How Esm-1 acts as a global mediator of glomerular gene expression (e.g., through epigenetic modifications, chemotaxis, etc.) will be an area for future study. A limitation of our study is that tools are still needed to tune the number of *Esm1*(+) cells or glomerular (vs. systemic) *Esm1* expression to assess the contribution of local Esm-1 to modulate global gene expression, podocyte number, and albuminuria.

The role of inflammation has been widely described in the development of DKD^15, 54–58^, and several studies outline a change in circulating Esm-1 concentrations in diabetes^59, 60^. Furthermore, Esm-1 is known as an anti-inflammatory proteoglycan^18, 19, 44^, and DKD susceptibility in mice has been linked to a deficiency in *Esm1* expression in glomerular endothelial cells^18, 41^. Esm-1 may significantly alter innate immunity genes as over-expression reduces albuminuria and alters expression of genes related to the interferon pathway^41^. In this study, over-expression of Esm-1 significantly altered expression of 18 interferon-related genes. Interestingly, 16 of these 18 genes were available in the bulk-RNA human database used in our study^33^, and 8 of these 16 interferon-related genes significantly correlate with *Esm1* expression, consistent with the findings reported by Zheng et al. Glomerular *Esm1* inversely correlated with glomerular macrophage infiltration; however, over-expression of Esm-1 did not reduce glomerular or tubulointerstitial macrophage infiltration^41^. Consistent with these findings, we also did not observe that Esm-1 over-expression appreciably reversed expression of genes that clustered with macrophage infiltration. Whether Esm-1 regulates a more specific subset of macrophages or whether only local Esm-1 modulates leukocyte infiltration is another area for further study.

In summary, our study shows that *Esm1* demonstrates glomerular expression in mice and humans. *Esm1* expression delineates a specialized subpopulation of mature endothelial cells, with a transcriptomic signature enriched for vascular and tissue development processes that changes with diabetes. Diabetes is associated with a fall in glomerular *Esm1* expression, with a transcriptional polarization characterized by variations of subsets of genes and pathways enriched in glomerular endothelial cells and in neighboring glomerular cells, tightly correlating with *Esm1* glomerular expression. Expression of genes enriched in glomerular compartments correlate with *Esm1* and albuminuria and can be rescued by systemic over-expression of Esm-1. From publicly available single cell and bulk transcriptomes, and through experimental validation by over-expression of Esm-1 in DKD-susceptible mice, our results generate novel mechanistic hypotheses about the role of *Esm1*, the subset of *Esm1*(+) cells, its role in glomerular crosstalk, and its role as sensor of transcriptional polarization in DKD.

## Supplementary Materials

### Supplementary Figure Legends

**Figure S1. Expression of *Esm1* compared to canonical genes of vasculature compartments in mouse kidney.**

(A-C) t-SNE plots of RECs, color-coded for the expression of *Esm1* (*purple*) and canonical genes of glomerular (g) (A), cortical (c) (B), and medullary (m) RECs (C). Magnitude of gene expression is indicated by the scale provided. (D) Violin plots showing the expression profiles of canonical genes of mouse kidney vasculature compartments in *Esm1*(+) and *Esm1*(-) RECs. (E-G) Heatmap of the relative fold-change in log-transformed expression-level of top 50 marker genes of subclusters in *Esm1*(+) vs. *Esm1*(-) glomerular (g) (E), cortical (c) (F), and medullary (m) RECs (G). All data are from single-cell mice databases M1, M2 and M3. RECs, renal endothelial cells.

**Figure S2. Expression of *Esm1* compared to canonical genes of vasculature compartments in human kidney.**

(A-E) t-SNE plots of RECs, color-coded for the expression of *Esm1* (*purple*) and canonical genes of afferent and efferent arterioles / peritubular capillaries (A), glomerular (B), and ascending vasa recta (C), descending vasa recta (D), and other endothelial cells (E). Magnitude of gene expression is indicated by the scale provided. (F) Violin plots showing the expression profiles of canonical genes of human kidney vasculature compartments in *Esm1*(+) and *Esm1*(-) RECs. All data are from single-cell human databases H1, H2 and H3. RECs, renal endothelial cells.

**Figure S3. Relationship between glomerular expression of *Esm1* and susceptibility to diabetic kidney disease in mice.**

(A) Quantification by RNAscope of *Esm1* glomerular expression in DBA/2 mice, compared to C57BL/6 mice. (B) Quantification by RNAscope of *Esm1* glomerular expression in DBA/2 mice with STZ-induced diabetic kidney disease, compared to controls. * p < 0.05, as indicated.

**Figure S4. Expression of *Esm1* in patients with diabetes.**

(A) t-SNE plots of RECs, color-coded for the expression of *Esm1* and canonical genes of human kidney vasculature compartments from patients with diabetes. Grey is no expression, light color is low expression, dark color is high expression. (B) Violin plots showing the expression profiles of canonical genes of human kidney vasculature compartments in *Esm1*(+) and *Esm1*(-) RECs from patients with diabetes. All data are from single-cell human databases H1 (control) and H4 (diabetes). RECs, renal endothelial cells.

**Figure S5. Glomerular RNA expression of genes co-expressed with *Esm1* at the single-cell level and from the Igfbp5 interactome.**

Z-scores of differential expressions of the top 15 differentially expressed genes on diverging branches according to trajectory inference analysis from patients with diabetes (A) and healthy controls (B). All data are from gene expression profile human databases H1 (control, *purple*) and H4 (diabetes, *green*).

**Figure S6. Glomerular RNA expression of genes co-expressed with *Esm1* at the single-cell level and from the Igfbp5 interactome.**

(A) Expression level of genes differentially expressed in *Esm1*(+) cells from healthy controls falling into the vascular development ontology cluster. We ordered genes from highest to lowest single cell expression fold-change in *Esm1*(+) vs. *Esm1*(-) cells. (B) Expression level of genes differentially expressed in *Esm1*(+) cells from healthy controls falling into the Chemotaxis ontology cluster. We ordered genes from highest to lowest single cell expression fold-change in *Esm1*(+) vs. *Esm1*(-) cells. (C) Expression level of genes from the Igfbp5 interactome falling into the extracellular matrix organization ontology. For each group, we displayed gene expression on the y-axis vs. *Esm1* expression on the x-axis and linear regression curves with 95% CI are shown. (E) Gene expression enriched in endothelial cells. (M) Gene expression enriched in mesangial cells. All data are from gene expression profile human databases H5 (control, *purple*) and H6 (diabetes, *green*).

**Figure S7. Overlaps between gene lists used in the human gene expression profile analysis.**

(A) Intersections between cell type-enriched whole gene lists used in the gene expression profile analysis. (B) Intersections between lists of cell type-enriched genes falling into the cluster of direct correlations and showing a significant correlation with *Esm1*. (C) Intersections between lists of cell type-enriched genes falling into the cluster of inverse correlations and showing a significant correlation with *Esm1*. All data are from gene expression profile the human database H6 (diabetes).

**Figure S8. Overlaps between gene lists used for the gene expression profile analysis in DBA/2 mice with streptozotocin-induced DKD.**

(A) Intersections between compartment-enriched whole gene lists used in the gene expression profile analysis. (B) Intersections between lists of compartment-enriched genes falling into the cluster of direct correlations with *Esm1* and inverse correlations with urine albumin-to-creatinine ratio (UACR). (C) Intersections between lists of compartment-enriched genes falling into the cluster of inverse correlations with *Esm1* and direct correlations with UACR. All data are from DBA/2 mice with streptozotocin-induced DKD (Database M4).

## Author Contributions

A.G., N.K., and V.B. designed the study; A.G., X.Z. and V.B. analyzed the data; A.G. and V.B. made the figures; A.G., X.Z. and V.B. drafted and revised the paper; all authors have seen and approved the final version of the manuscript.

## Acknowledgments

A.G. was funded by a Fulbright Hauts-de-France grant and by the France – Stanford Center for Interdisciplinary studies. X.Z. was funded in part by a Larry L. Hillblom Foundation Postdoctoral Fellowship award and the Holmgren Family Foundation. V.B. was funded, in part, by the NIDDK (R01 DK091565).

The authors thank Drs. Rajasree Menon and Matthias Kretzler for sharing of data information.

## Abbreviations

AEA: afferent and efferent arterioles
AVR: ascending vasa recta
cRECs: cortical renal endothelial cells
DKD: diabetic kidney disease
DVR: descending vasa recta
Esm-1: endothelial cell-specific molecule-1
gRECs: glomerular renal endothelial cells
mRECs: medullary renal endothelial cells
PTC: peritubular capillaries

## References

1. Parving HH, Hommel E, Mathiesen E, Skøtt P, Edsberg B, Bahnsen M, et al. Prevalence of microalbuminuria, arterial hypertension, retinopathy and neuropathy in patients with insulin dependent diabetes. Br Med J Clin Res Ed. 1988 Jan 16;296(6616):156–60.

2. Borch-Johnsen K, Andersen PK, Deckert T. The effect of proteinuria on relative mortality in type 1 (insulin-dependent) diabetes mellitus. Diabetologia. 1985 Aug;28(8):590–6.

3. Rossing P, Hougaard P, Borch-Johnsen K, Parving HH. Predictors of mortality in insulin dependent diabetes: 10 year observational follow up study. BMJ. 1996 Sep 28;313(7060):779–84.

4. Gurley SB, Clare SE, Snow KP, Hu A, Meyer TW, Coffman TM. Impact of genetic background on nephropathy in diabetic mice. Am J Physiol Renal Physiol. 2006 Jan;290(1):F214–222.

5. Qi Z, Fujita H, Jin J, Davis LS, Wang Y, Fogo AB, et al. Characterization of susceptibility of inbred mouse strains to diabetic nephropathy. Diabetes. 2005 Sep;54(9):2628–37.

6. Breyer MD, Böttinger E, Brosius FC, Coffman TM, Harris RC, Heilig CW, et al. Mouse models of diabetic nephropathy. J Am Soc Nephrol JASN. 2005 Jan;16(1):27–45.

7. Chow FY, Nikolic-Paterson DJ, Ma FY, Ozols E, Rollins BJ, Tesch GH. Monocyte chemoattractant protein-1-induced tissue inflammation is critical for the development of renal injury but not type 2 diabetes in obese db/db mice. Diabetologia. 2007 Feb;50(2):471–80.

8. Chow FY, Nikolic-Paterson DJ, Ozols E, Atkins RC, Tesch GH. Intercellular adhesion molecule-1 deficiency is protective against nephropathy in type 2 diabetic db/db mice. J Am Soc Nephrol JASN. 2005 Jun;16(6):1711–22.

9. Okada S, Shikata K, Matsuda M, Ogawa D, Usui H, Kido Y, et al. Intercellular adhesion molecule-1-deficient mice are resistant against renal injury after induction of diabetes. Diabetes. 2003 Oct;52(10):2586–93.

10. Bhalla V. The Role of the Immune System in the Pathogenesis of Diabetic Nephropathy. J Nephrol Ther. 2012 Jan 1;s2.

11. Moon JY, Jeong KH, Lee TW, Ihm CG, Lim SJ, Lee SH. Aberrant recruitment and activation of T cells in diabetic nephropathy. Am J Nephrol. 2012;35(2):164–74.

12. Hickey FB, Martin F. Role of the Immune System in Diabetic Kidney Disease. Curr Diab Rep. 2018 12;18(4):20.

13. Awad AS, You H, Gao T, Cooper TK, Nedospasov SA, Vacher J, et al. Macrophage-derived tumor necrosis factor-α mediates diabetic renal injury. Kidney Int. 2015 Oct;88(4):722–33.

14. Chow FY, Nikolic-Paterson DJ, Atkins RC, Tesch GH. Macrophages in streptozotocin-induced diabetic nephropathy: potential role in renal fibrosis. Nephrol Dial Transplant Off Publ Eur Dial Transpl Assoc - Eur Ren Assoc. 2004 Dec;19(12):2987–96.

15. Nguyen D, Ping F, Mu W, Hill P, Atkins RC, Chadban SJ. Macrophage accumulation in human progressive diabetic nephropathy. Nephrol Carlton Vic. 2006 Jun;11(3):226–31.

16. Lim AKH, Ma FY, Nikolic-Paterson DJ, Kitching AR, Thomas MC, Tesch GH. Lymphocytes promote albuminuria, but not renal dysfunction or histological damage in a mouse model of diabetic renal injury. Diabetologia. 2010 Aug;53(8):1772–82.

17. Lassalle P, Molet S, Janin A, Heyden JV, Tavernier J, Fiers W, et al. ESM-1 is a novel human endothelial cell-specific molecule expressed in lung and regulated by cytokines. J Biol Chem. 1996 Aug 23;271(34):20458–64.

18. Zheng X, Soroush F, Long J, Hall ET, Adishesha PK, Bhattacharya S, et al. Murine glomerular transcriptome links endothelial cell-specific molecule-1 deficiency with susceptibility to diabetic nephropathy. PloS One. 2017;12(9):e0185250.

19. Béchard D, Scherpereel A, Hammad H, Gentina T, Tsicopoulos A, Aumercier M, et al. Human endothelial-cell specific molecule-1 binds directly to the integrin CD11a/CD18 (LFA-1) and blocks binding to intercellular adhesion molecule-1. J Immunol Baltim Md 1950. 2001 Sep 15;167(6):3099–106.

20. De Freitas Caires N, Gaudet A, Portier L, Tsicopoulos A, Mathieu D, Lassalle P. Endocan, sepsis, pneumonia, and acute respiratory distress syndrome. Crit Care Lond Engl. 2018 Oct 26;22(1):280.

21. Zheng X, Higdon L, Gaudet A, Shah M, Balistieri A, Li C, et al. Endothelial Cell-Specific Molecule-1 Inhibits Albuminuria in Diabetic Mice [Internet]. 2021 Oct [cited 2021 Dec 20] p. 2021.10.14.464296. Available from: https://www.biorxiv.org/content/10.1101/2021.10.14.464296v1

22. Rocha SF, Schiller M, Jing D, Li H, Butz S, Vestweber D, et al. Esm1 modulates endothelial tip cell behavior and vascular permeability by enhancing VEGF bioavailability. Circ Res. 2014 Aug 29;115(6):581–90.

23. Pontes-Quero S, Fernández-Chacón M, Luo W, Lunella FF, Casquero-Garcia V, Garcia-Gonzalez I, et al. High mitogenic stimulation arrests angiogenesis. Nat Commun. 2019 01;10(1):2016.

24. del Toro R, Prahst C, Mathivet T, Siegfried G, Kaminker JS, Larrivee B, et al. Identification and functional analysis of endothelial tip cell-enriched genes. Blood. 2010 Nov 11;116(19):4025–33.

25. Yang H, Fang L, Zhan R, Hegarty JM, Ren J, Hsiai TK, et al. Polo-like kinase 2 regulates angiogenic sprouting and blood vessel development. Dev Biol. 2015 Aug 15;404(2):49–60.

26. Barry DM, McMillan EA, Kunar B, Lis R, Zhang T, Lu T, et al. Molecular determinants of nephron vascular specialization in the kidney. Nat Commun. 2019 13;10(1):5705.

27. Dumas SJ, Meta E, Borri M, Goveia J, Rohlenova K, Conchinha NV, et al. Single-Cell RNA Sequencing Reveals Renal Endothelium Heterogeneity and Metabolic Adaptation to Water Deprivation. J Am Soc Nephrol JASN. 2020 Jan;31(1):118–38.

28. Ransick A, Lindström NO, Liu J, Zhu Q, Guo JJ, Alvarado GF, et al. Single-Cell Profiling Reveals Sex, Lineage, and Regional Diversity in the Mouse Kidney. Dev Cell. 2019 Nov 4;51(3):399–413.e7.

29. Wilson PC, Wu H, Kirita Y, Uchimura K, Ledru N, Rennke HG, et al. The single-cell transcriptomic landscape of early human diabetic nephropathy. Proc Natl Acad Sci U S A. 2019 24;116(39):19619–25.

30. Lake BB, Chen S, Hoshi M, Plongthongkum N, Salamon D, Knoten A, et al. A single-nucleus RNA-sequencing pipeline to decipher the molecular anatomy and pathophysiology of human kidneys. Nat Commun. 2019 27;10(1):2832.

31. Stewart BJ, Ferdinand JR, Young MD, Mitchell TJ, Loudon KW, Riding AM, et al. Spatiotemporal immune zonation of the human kidney. Science. 2019 27;365(6460):1461–6.

32. Menon R, Otto EA, Hoover P, Eddy S, Mariani L, Godfrey B, et al. Single cell transcriptomics identifies focal segmental glomerulosclerosis remission endothelial biomarker. JCI Insight. 2020 Mar 26;5(6).

33. Pan Y, Jiang S, Hou Q, Qiu D, Shi J, Wang L, et al. Dissection of Glomerular Transcriptional Profile in Patients With Diabetic Nephropathy: SRGAP2a Protects Podocyte Structure and Function. Diabetes. 2018;67(4):717–30.

34. Chen H, Albergante L, Hsu JY, Lareau CA, Lo Bosco G, Guan J, et al. Single-cell trajectories reconstruction, exploration and mapping of omics data with STREAM. Nat Commun. 2019 23;10(1):1903.

35. Guo G, Pinello L, Han X, Lai S, Shen L, Lin TW, et al. Serum-Based Culture Conditions Provoke Gene Expression Variability in Mouse Embryonic Stem Cells as Revealed by Single-Cell Analysis. Cell Rep. 2016 Feb 2;14(4):956–65.

36. Zhang Z, Wang J. MLLE: Modified Locally Linear Embedding Using Multiple Weights. In: Schölkopf B, Platt JC, Hoffman T, editors. Advances in Neural Information Processing Systems 19 [Internet]. MIT Press; 2007 [cited 2020 Apr 9]. p. 1593–600. Available from: http://papers.nips.cc/paper/3132-mlle-modified-locally-linear-embedding-using-multiple-weights.pdf

37. Park J, Shrestha R, Qiu C, Kondo A, Huang S, Werth M, et al. Single-cell transcriptomics of the mouse kidney reveals potential cellular targets of kidney disease. Science. 2018 18;360(6390):758–63.

38. Karaiskos N, Rahmatollahi M, Boltengagen A, Liu H, Hoehne M, Rinschen M, et al. A Single-Cell Transcriptome Atlas of the Mouse Glomerulus. J Am Soc Nephrol JASN. 2018;29(8):2060–8.

39. Zhou Y, Zhou B, Pache L, Chang M, Khodabakhshi AH, Tanaseichuk O, et al. Metascape provides a biologist-oriented resource for the analysis of systems-level datasets. Nat Commun. 2019 03;10(1):1523.

40. Lex A, Gehlenborg N, Strobelt H, Vuillemot R, Pfister H. UpSet: Visualization of Intersecting Sets. IEEE Trans Vis Comput Graph. 2014 Dec;20(12):1983–92.

41. Zheng X, Higdon L, Gaudet A, Shah M, Balistieri A, Li C, et al. Endothelial Cell-Specific Molecule-1 Inhibits Albuminuria in Diabetic Mice. Kidney360. 2022 Dec 29;3(12):2059–76.

42. S Y, X C, X C, M T. Analysing the meta-interaction between pathways by gene set topological impact analysis. BMC Genomics [Internet]. 2020 Oct 27 [cited 2023 Feb 22];21(1). Available from: https://pubmed.ncbi.nlm.nih.gov/33109101/

43. Huang J, Kong Y, Xie C, Zhou L. Stem/progenitor cell in kidney: characteristics, homing, coordination, and maintenance. Stem Cell Res Ther. 2021 Mar 20;12(1):197.

44. Gaudet A, Portier L, Prin M, Copin MC, Tsicopoulos A, Mathieu D, et al. Endocan regulates acute lung inflammation through control of leukocyte diapedesis. J Appl Physiol Bethesda Md 1985. 2019 Sep 1;127(3):668–78.

45. Zhang X, Zhuang R, Wu H, Chen J, Wang F, Li G, et al. A novel role of endocan in alleviating LPS-induced acute lung injury. Life Sci. 2018 Jun 1;202:89–97.

46. Sison K, Eremina V, Baelde H, Min W, Hirashima M, Fantus IG, et al. Glomerular structure and function require paracrine, not autocrine, VEGF-VEGFR-2 signaling. J Am Soc Nephrol JASN. 2010 Oct;21(10):1691–701.

47. Jeansson M, Gawlik A, Anderson G, Li C, Kerjaschki D, Henkelman M, et al. Angiopoietin-1 is essential in mouse vasculature during development and in response to injury. J Clin Invest. 2011 Jun;121(6):2278–89.

48. Vaughan MR, Quaggin SE. How do mesangial and endothelial cells form the glomerular tuft? J Am Soc Nephrol JASN. 2008 Jan;19(1):24–33.

49. Schlöndorff D, Banas B. The mesangial cell revisited: no cell is an island. J Am Soc Nephrol JASN. 2009 Jun;20(6):1179–87.

50. Nam TJ, Busby WH, Rees C, Clemmons DR. Thrombospondin and osteopontin bind to insulin-like growth factor (IGF)-binding protein-5 leading to an alteration in IGF-I-stimulated cell growth. Endocrinology. 2000 Mar;141(3):1100–6.

51. Gui Y, Murphy LJ. Interaction of insulin-like growth factor binding protein-3 with latent transforming growth factor-beta binding protein-1. Mol Cell Biochem. 2003 Aug;250(1– 2):189–95.

52. Loechel F, Fox JW, Murphy G, Albrechtsen R, Wewer UM. ADAM 12-S cleaves IGFBP-3 and IGFBP-5 and is inhibited by TIMP-3. Biochem Biophys Res Commun. 2000 Nov 30;278(3):511–5.

53. Amaar YG, Thompson GR, Linkhart TA, Chen ST, Baylink DJ, Mohan S. Insulin-like growth factor-binding protein 5 (IGFBP-5) interacts with a four and a half LIM protein 2 (FHL2). J Biol Chem. 2002 Apr 5;277(14):12053–60.

54. Galkina E, Ley K. Leukocyte recruitment and vascular injury in diabetic nephropathy. J Am Soc Nephrol JASN. 2006 Feb;17(2):368–77.

55. Tesch GH. Macrophages and diabetic nephropathy. Semin Nephrol. 2010 May;30(3):290–301.

56. Gohda T, Niewczas MA, Ficociello LH, Walker WH, Skupien J, Rosetti F, et al. Circulating TNF receptors 1 and 2 predict stage 3 CKD in type 1 diabetes. J Am Soc Nephrol JASN. 2012 Mar;23(3):516–24.

57. Verzola D, Cappuccino L, D’Amato E, Villaggio B, Gianiorio F, Mij M, et al. Enhanced glomerular Toll-like receptor 4 expression and signaling in patients with type 2 diabetic nephropathy and microalbuminuria. Kidney Int. 2014 Dec;86(6):1229–43.

58. Zhang C, Xiao C, Wang P, Xu W, Zhang A, Li Q, et al. The alteration of Th1/Th2/Th17/Treg paradigm in patients with type 2 diabetes mellitus: Relationship with diabetic nephropathy. Hum Immunol. 2014 Apr;75(4):289–96.

59. Cikrikcioglu MA, Erturk Z, Kilic E, Celik K, Ekinci I, Yasin Cetin AI, et al. Endocan and albuminuria in type 2 diabetes mellitus. Ren Fail. 2016 Nov;38(10):1647–53.

60. Ekiz-Bilir B, Bilir B, Aydın M, Soysal-Atile N. Evaluation of endocan and endoglin levels in chronic kidney disease due to diabetes mellitus. Arch Med Sci AMS. 2019 Jan;15(1):86–91.

