## Supplementary figures and images for "Esm-1 mediates transcriptional polarization associated with diabetic kidney disease"

### Figure S1

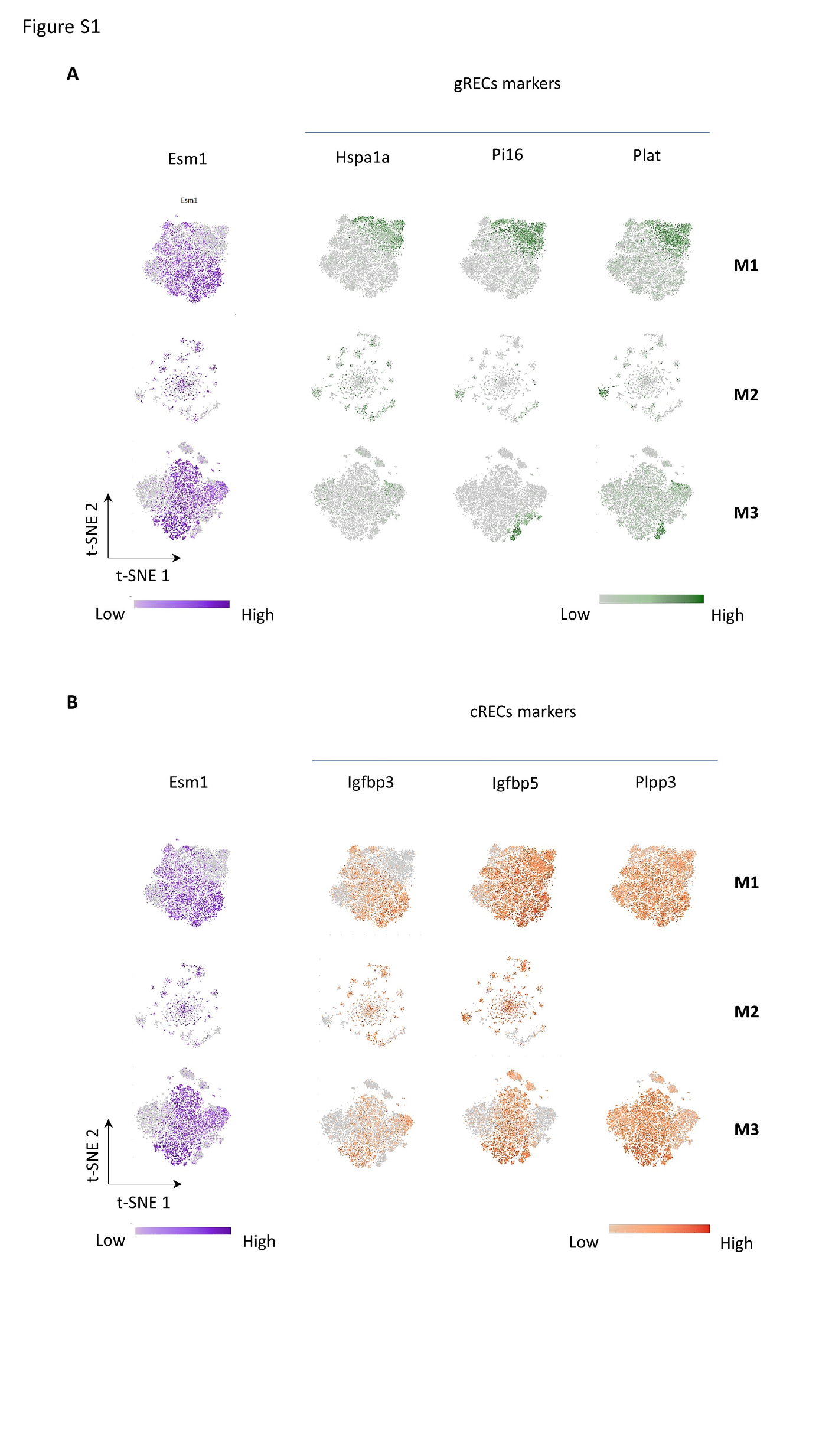

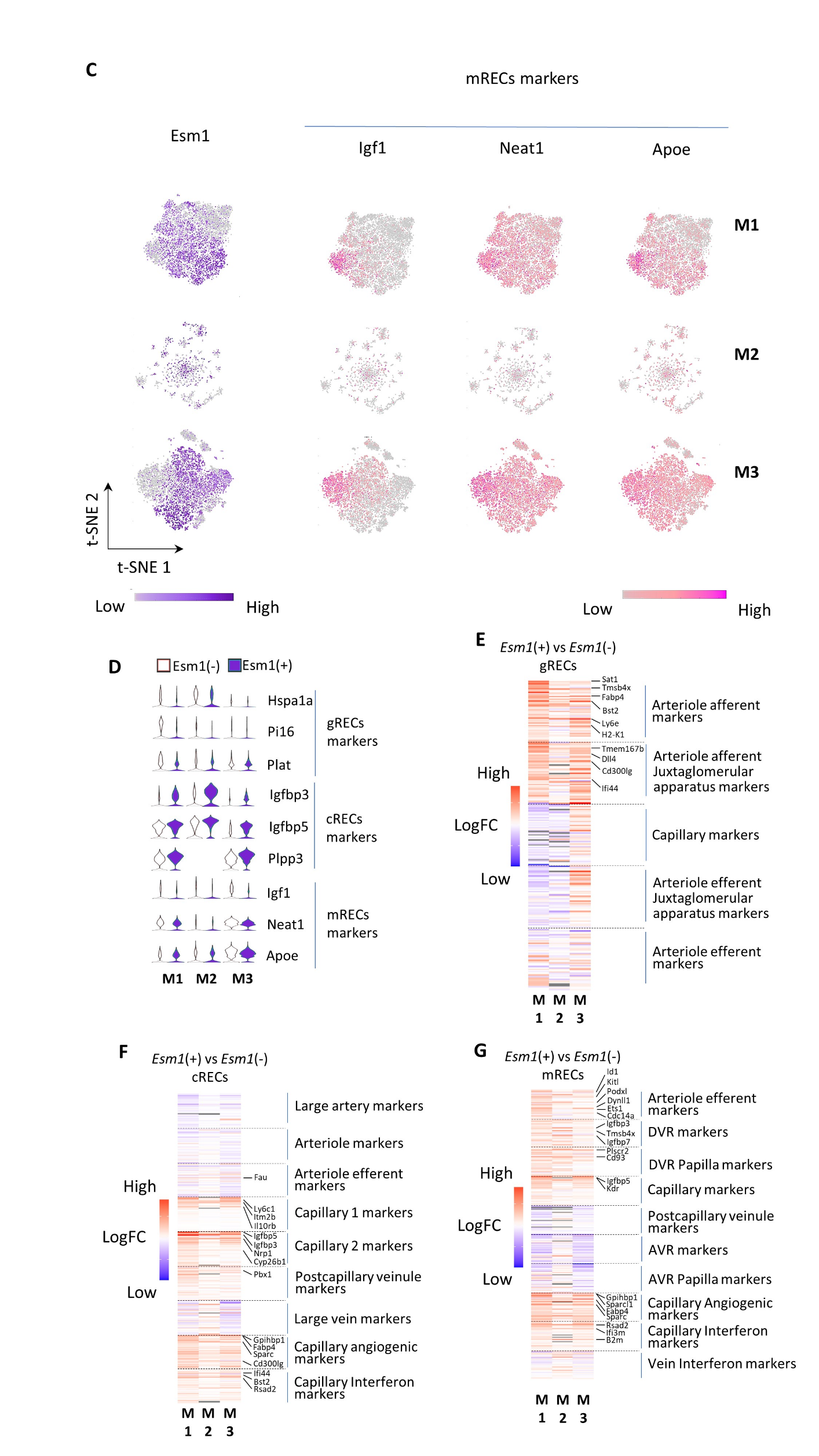

### Figure S2

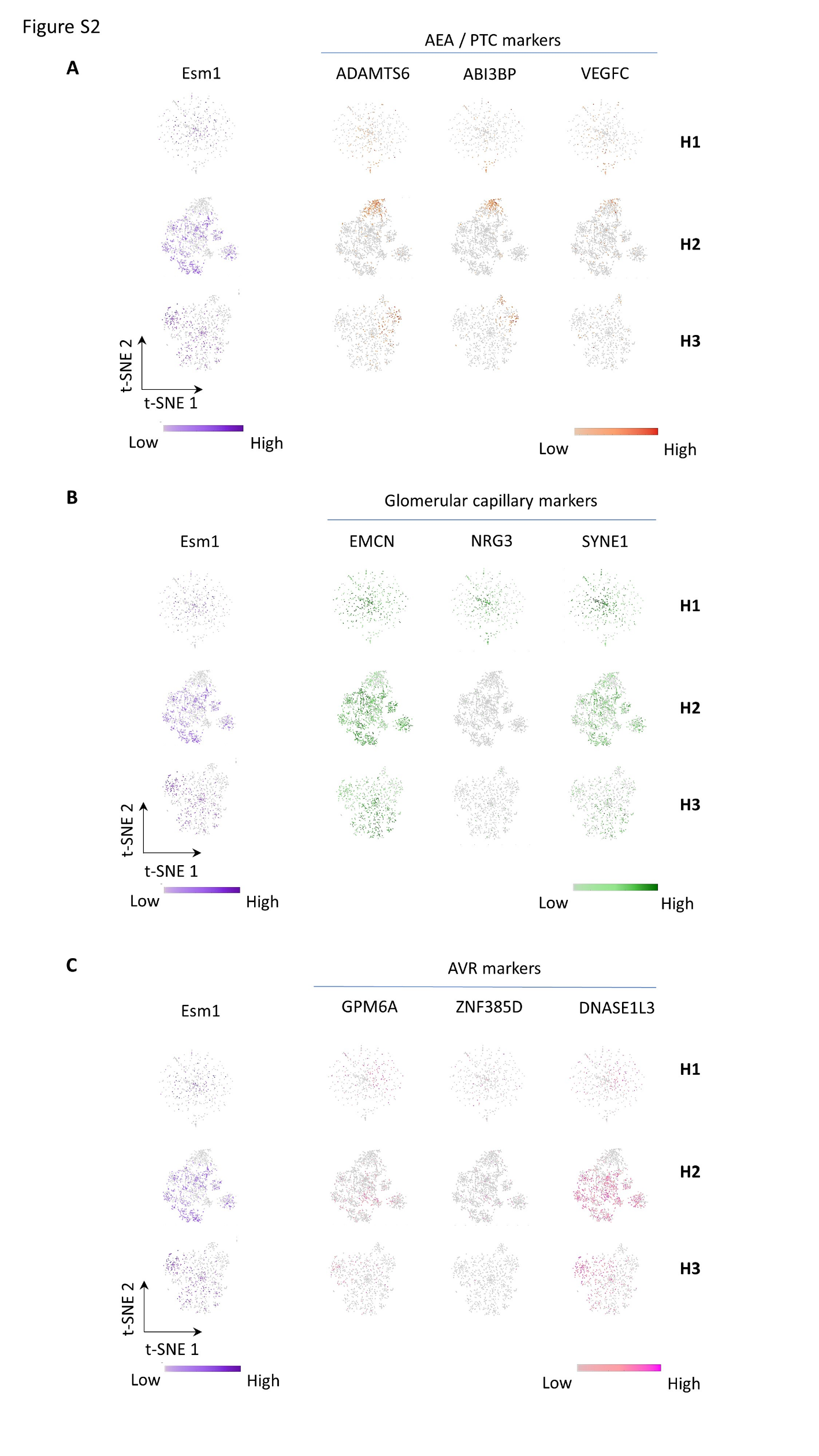

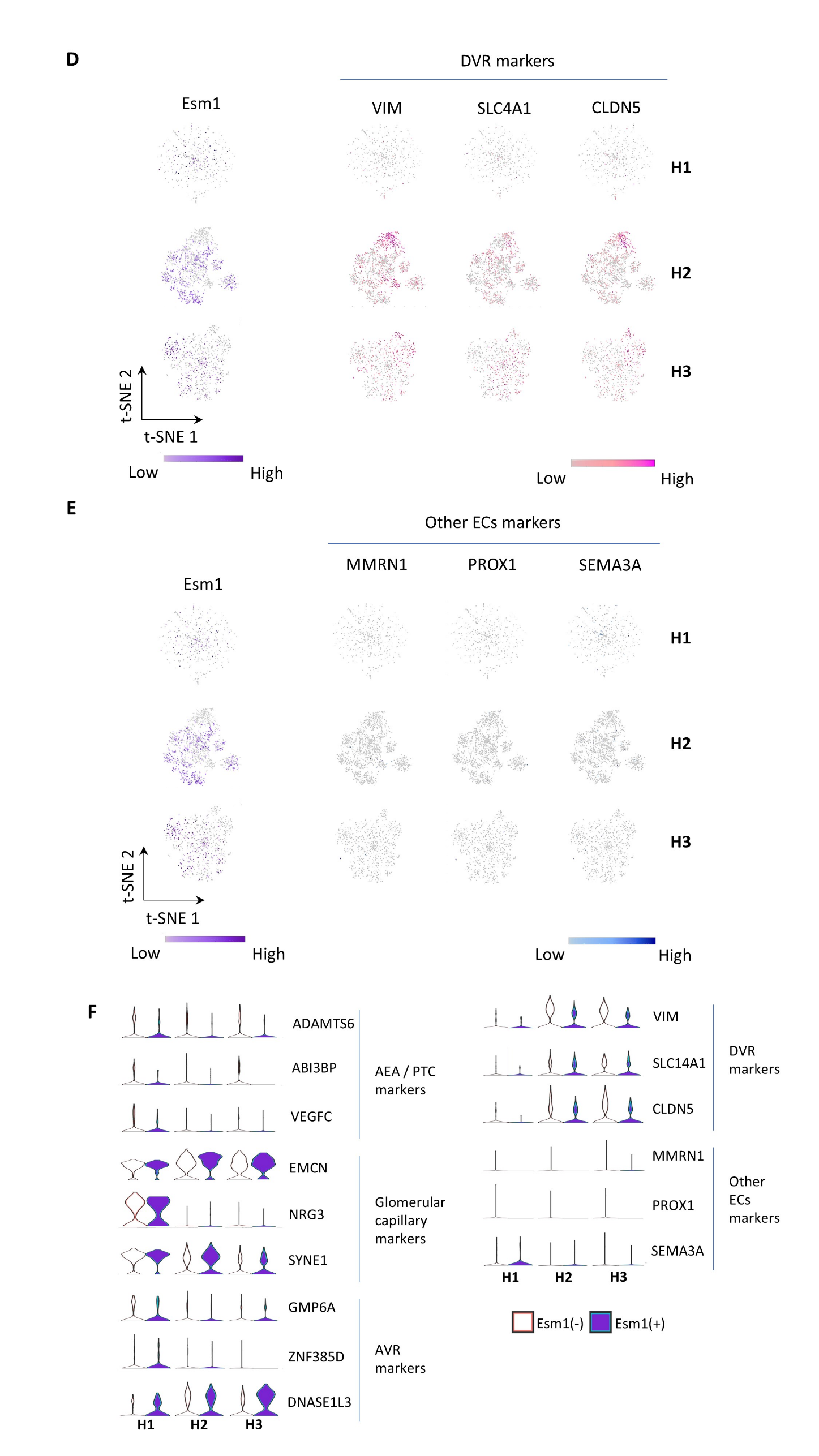

### Figure S3

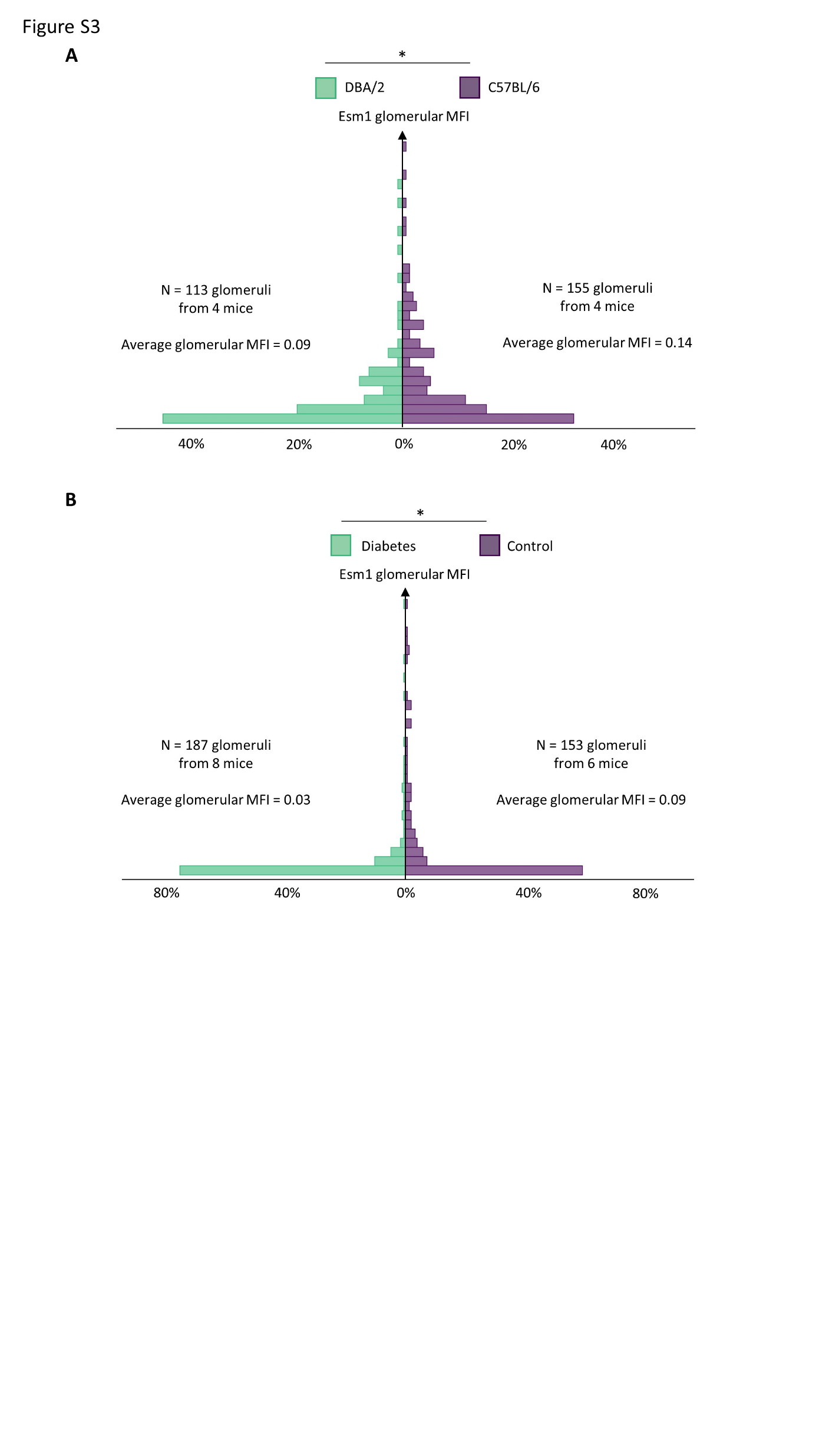

### Figure S4

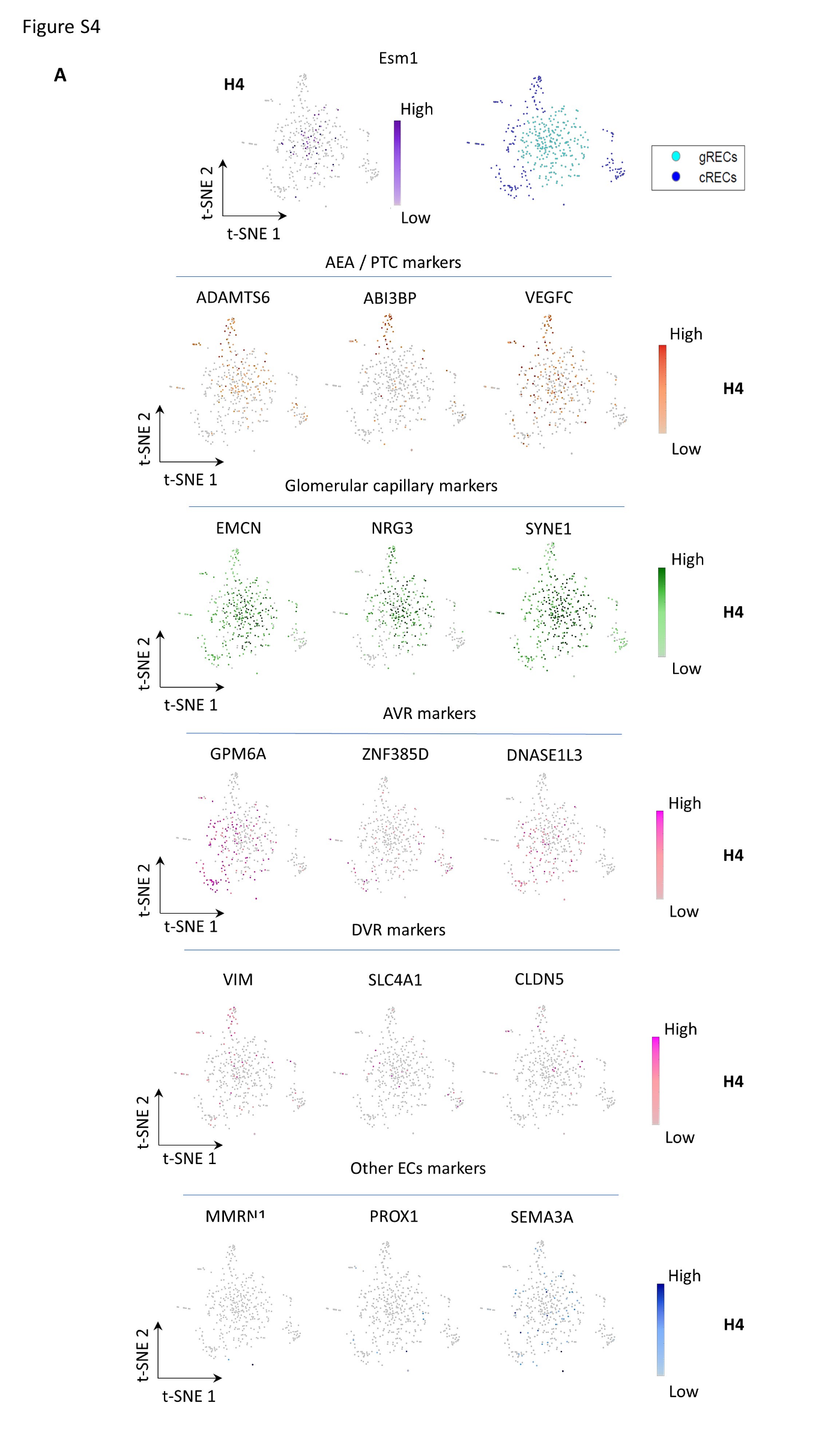

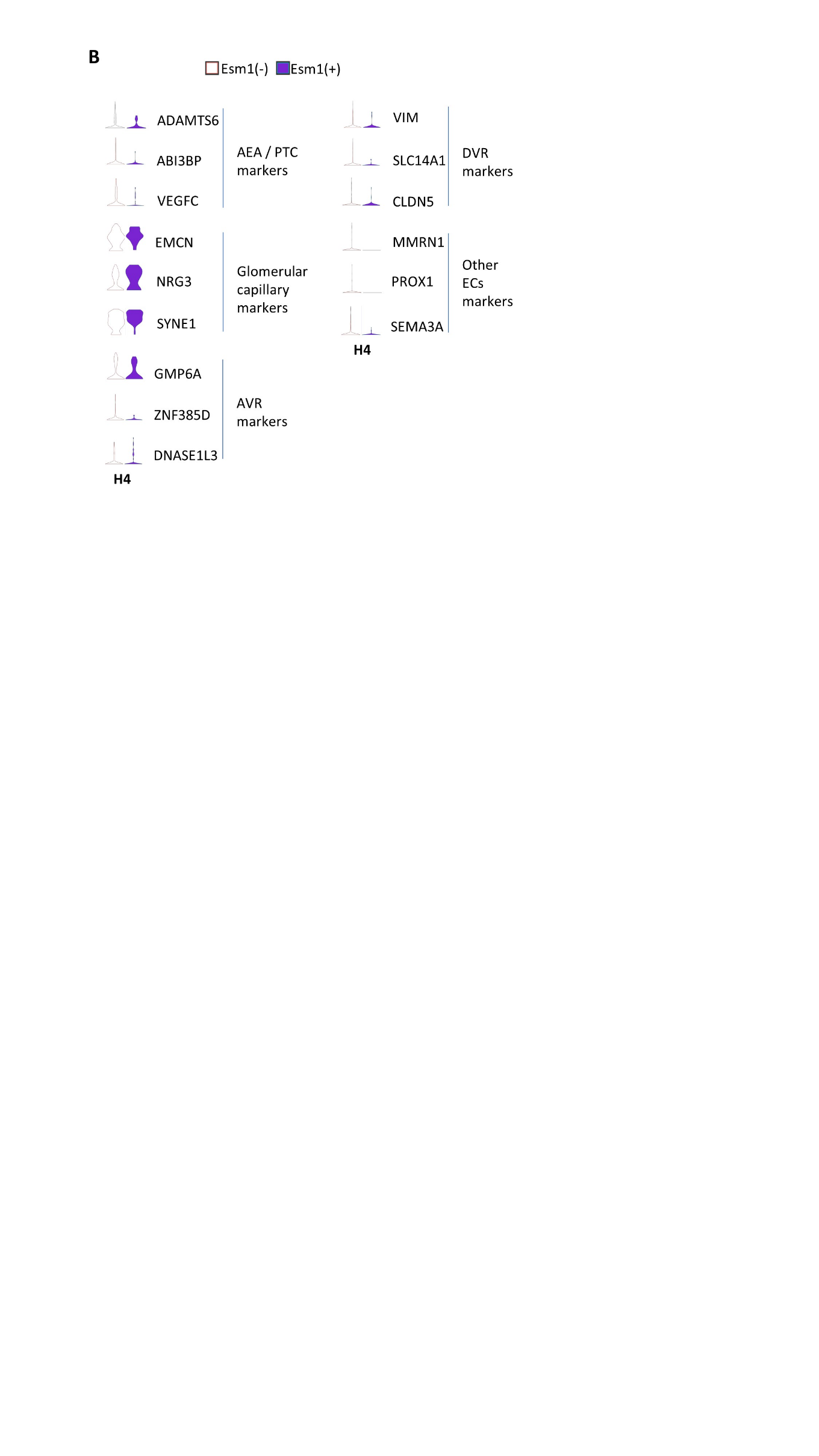

### Figure S5

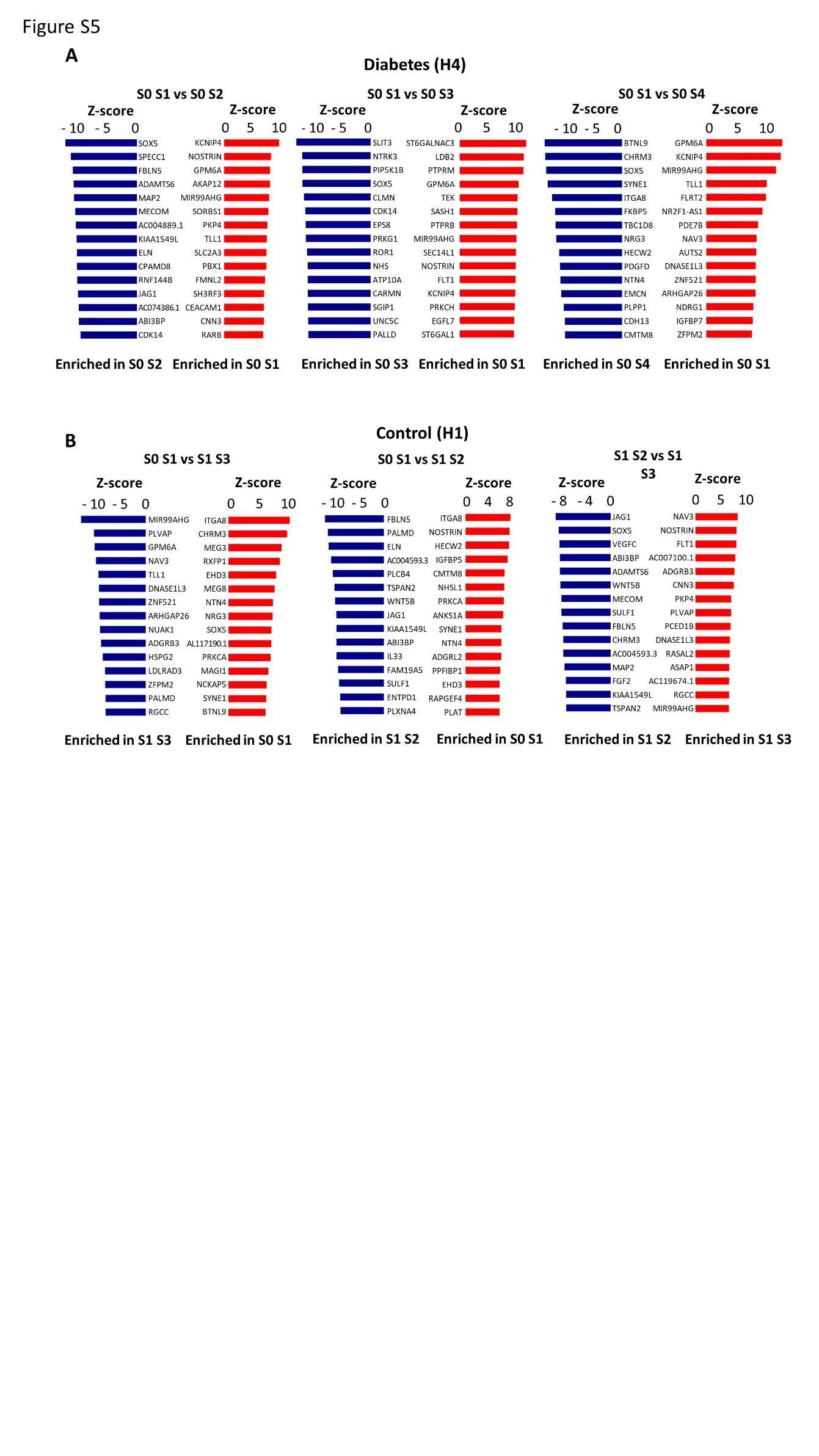

### Figure S6

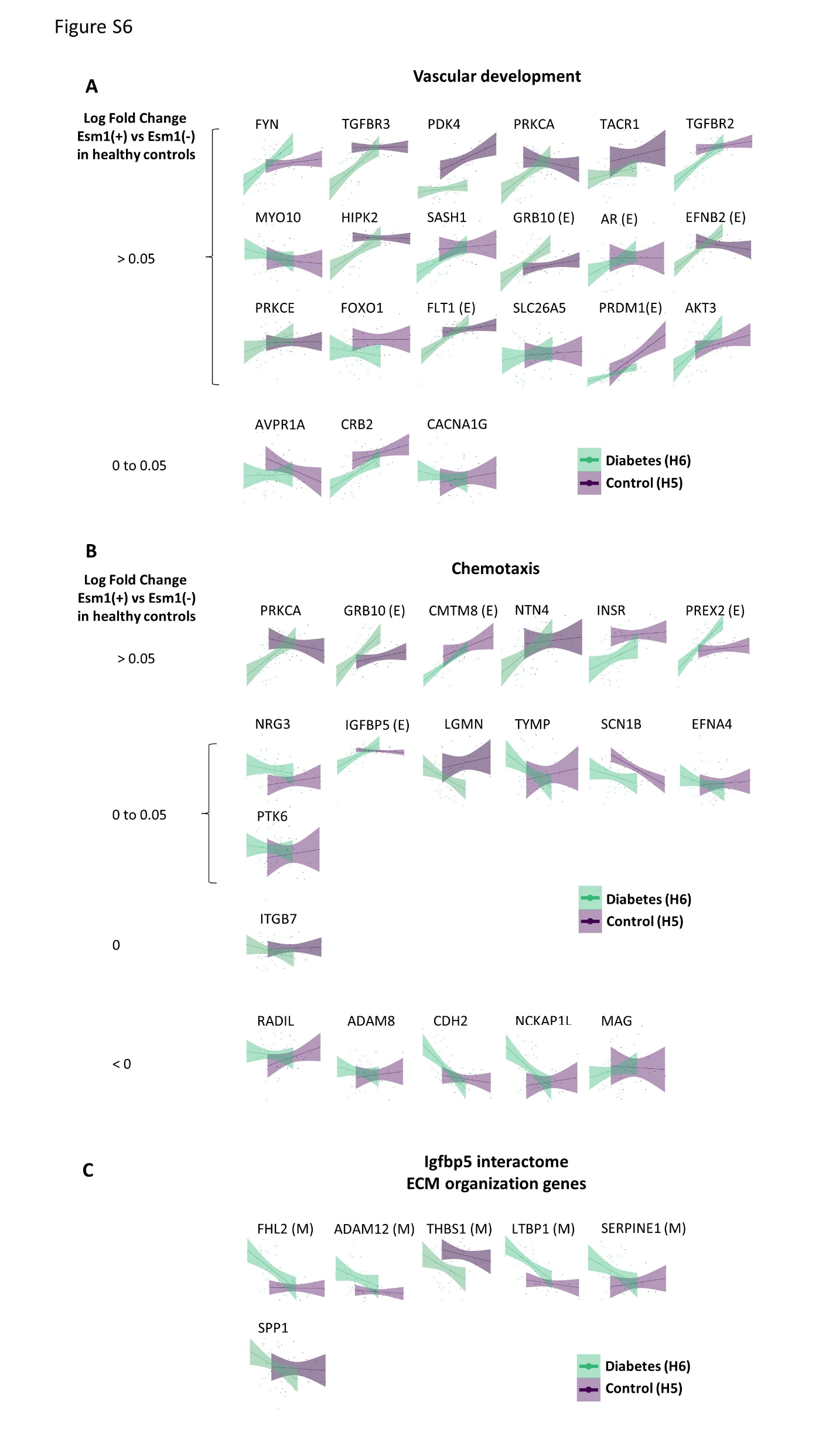

### Figure S7

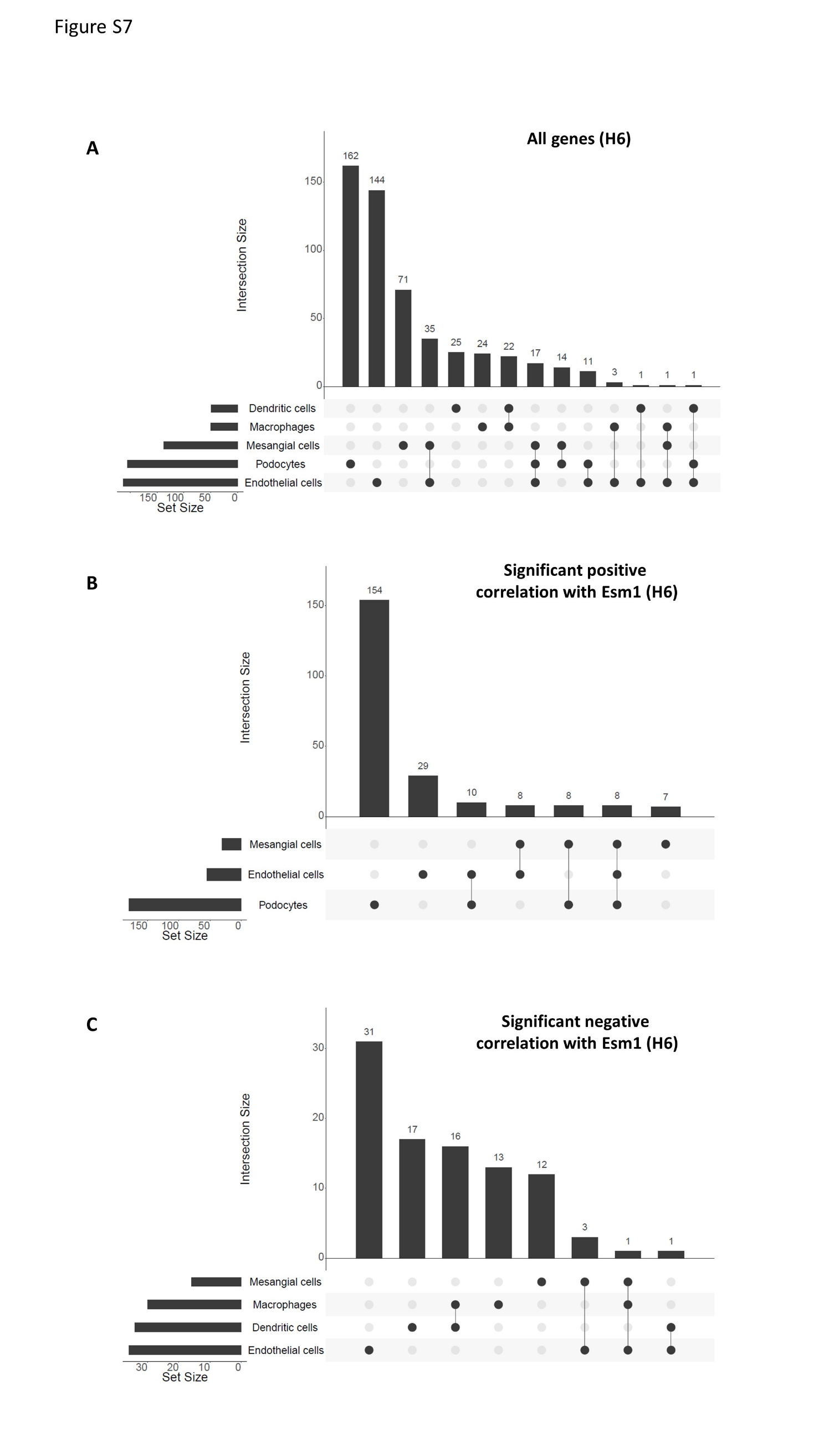

### Figure S8

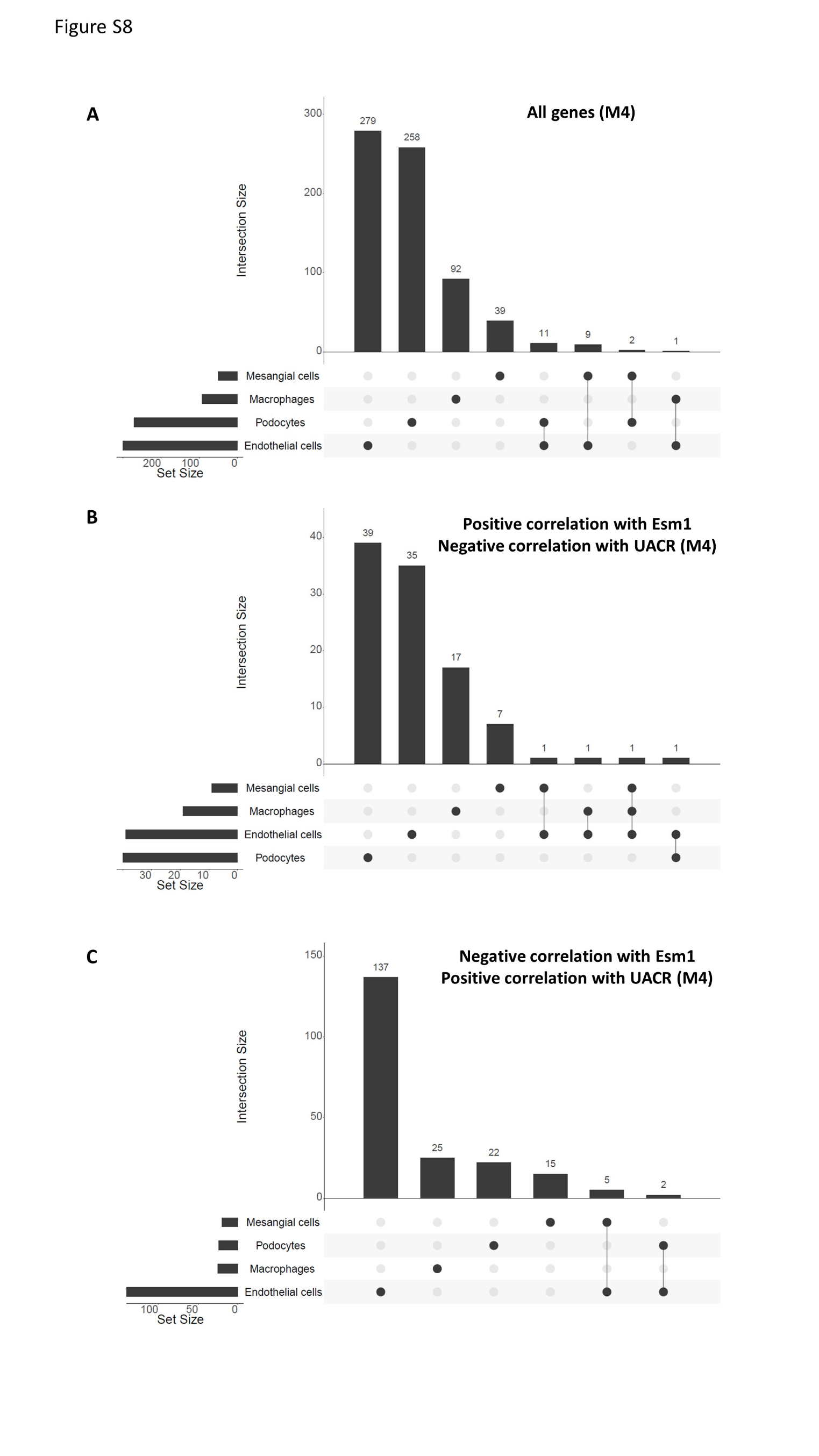
